# Decoding the *BRCA2* reversion principles underlying PARP inhibitor resistance

**DOI:** 10.64898/2026.04.29.721733

**Authors:** Anna Gabrielle Horacek, Frances Li Kueper, Robert Lu, Carter Joseph Simington, Isabella Paige Lamont, Stella Tran, Hanqin Li, Dirk Hockemeyer

## Abstract

Reversion mutations that restore BRCA2 function represent a major mechanism of resistance to poly(ADP-ribose) polymerase inhibitors (PARPi) in *BRCA2*-mutant cancers. Predicting these events could inform treatment strategies, identify patients at increased risk of acquiring PARPi resistance, and improve interpretation of secondary variants. Here, we use an isogenic cell system and CRISPR editing to define the principles governing *BRCA2* reversion. We show that local sequence context shapes the spectrum of reading-frame restoration events, whereas domain architecture determines which of these events are selected as PARPi-resistant reversions. We characterize two routes of *BRCA2* restoration that operate both within and across exons: DNA-level reading-frame restoration and transcript-level rescue via alternative splicing. Lastly, we identify distinct exon 11 restoration mechanisms, including large genomic deletions and recurrent splice isoforms that can bypass the loss of the canonical splice donor. These findings provide a basis for defining a predictive code for *BRCA2* reversions that confer PARPi resistance.

## Introduction

Approximately 5% of breast cancers carry biallelic mutations in the BRCA1 or BRCA2 tumor suppressor proteins^1–3^. BRCA1/2 are critical for the high-fidelity repair of DNA double-strand breaks (DSBs) by facilitating homologous recombination (HR)^4,5^. BRCA2 contains several well-characterized functional domains, including the PALB2 binding domain^6^ and the RAD51-interacting BRC repeats^7^. In BRCA2 loss-of-function (LOF) cells, treatment with a PARP inhibitor (PARPi) induces excessive DNA damage and cell death, a phenomenon known as synthetic lethality^8–12^. PARPi sensitivity has been assessed in BRCA2 LOF tumors using RAD51 nuclear foci formation as a functional readout for HR activity^13^. Around 40-50% of patients with BRCA1/2 negative cancers do not respond to PARPi initially, and the majority of patients who do respond eventually acquire resistance^14^. A common form of acquired PARPi resistance arises from genomic reversion of the original *BRCA2* loss-of-function (LOF) mutation, in which a secondary somatic mutation restores the *BRCA2* reading frame and HR capacity^15,16^.

Secondary mutations can restore an in-frame *BRCA2* coding sequence through at least two distinct mechanisms^17^. The first restores the *BRCA2* reading frame at the DNA level through the genetic restoration of the wild-type sequence, second-site deletions or insertions, or large in-frame deletions that encompass the original mutation^17^. The second restores the *BRCA2* reading frame at the transcript level through splicing-mediated restoration, producing in-frame isoforms that bypass the pathogenic mutation or restore the reading frame^18^. Not all molecular events that restore an in-frame *BRCA2* coding sequence necessarily restore BRCA2 function or confer PARPi resistance. We therefore distinguish between sequence restoration and functional reversion. In this study, we use the term restoration event for any molecular event that re-establishes an in-frame *BRCA2* coding sequence, and reserve reversion for restoration events that restore BRCA2 function, fully or in part, and confer PARPi resistance. This distinction is particularly important given the positional bias observed among clinical *BRCA2* reversions. Although pathogenic mutations occur at similar frequencies across *BRCA2*, reversions are unevenly distributed. Clinical reversions occur more frequently in the N-terminal “hot spot” (c.750– 775), whereas reversions are less frequent in the C-terminal “cold spot” (after c.7617)^17^. Collectively, these observations point to an underlying “reversion code” that governs the likelihood that an initial BRCA2 mutation will acquire a secondary mutation capable of conferring PARPi resistance.

A key limitation in deciphering this reversion code is that studies have primarily relied on retrospective analyses of patient samples and on individual loci in cell culture systems^15,17,19,20^. While informative, these approaches generally capture only the spectrum of reversion mutations detectable after treatment selection and may therefore underappreciate secondary mutations that restore the reading frame but fail to confer resistance. Here, we use *BRCA2* locally haploid (loHAP) cell lines to systematically annotate restoration events across the *BRCA2* coding sequence and determine which confer PARPi resistance in an isogenic background^21^. *BRCA2* loHAPs are engineered by removing one copy of the *BRCA2* gene using CRISPR/Cas9 genome editing, thereby creating a haploid state in which mutations can be evaluated with high-throughput next-generation sequencing (NGS). Using CRISPR-Cas9 editing, lentivirus-based CRISPR screening, and long-read nanopore sequencing, we characterized the landscape of *BRCA2* reversions. Together, our results suggest a *BRCA2* “reversion code” in which local sequence features and regional protein context shape both the formation of restoration events and selection of PARPi-resistant reversions, and we further characterize the DNA- and transcript-associated mechanisms through which these events occur.

## Results

### *BRCA2* frameshift mutations induce PARPi sensitivity

To establish a systematic platform for evaluating reversion mutations, we generated a panel of isogenic *BRCA2* frameshift mutations and assayed their sensitivity to PARPi. Frameshift mutations were introduced in exons 2, 3, 5, 11, 12, 15, 23, and 27 using CRISPR-Cas9 editing with a single sgRNA in a *BRCA2* loHAP cell line (Fig. 1A)^21^. This loHAP cell line was derived from WIBR3 human embryonic stem cells (hESCs) and has a single copy of *BRCA2*, as confirmed by array comparative genomic hybridization^21,22^. Although loss-of-function frameshift alleles are outcompeted by unedited *BRCA2* loHAP cells in pooled culture, clonal frameshift cell lines can be isolated when cells are arrayed immediately after CRISPR-Cas9 editing. To test their PARPi sensitivity, clonal frameshift cell lines were mixed with unedited *BRCA2* loHAP cells at a 1:1 (50:50) ratio and treated with mock medium, 0.5 μM niraparib, or 1 μM niraparib for five days. Relative allele frequencies were then measured by NGS of the region surrounding the primary frameshift mutation. Frameshift alleles were further depleted relative to unedited cells even under mock conditions, consistent with a documented proliferative disadvantage following loss of BRCA2 function^21,23^. Frameshift alleles were also strongly depleted under niraparib treatment, with significant depletion occurring at 0.5 μM niraparib for all exons tested (Fig. 1B, C). The degree of depletion was highly consistent for exons 2 - 11 and 23. By contrast, frameshift mutations in exon 12 (p.L2304Ffs*34) and exon 27 (p.S3291Ffs*34) were less sensitive to PARPi treatment, suggesting that these alleles retain partial BRCA2 function, consistent with prior reports that exon 27 can produce hypomorphic phenotypes^24,25^.

**Figure 1:**
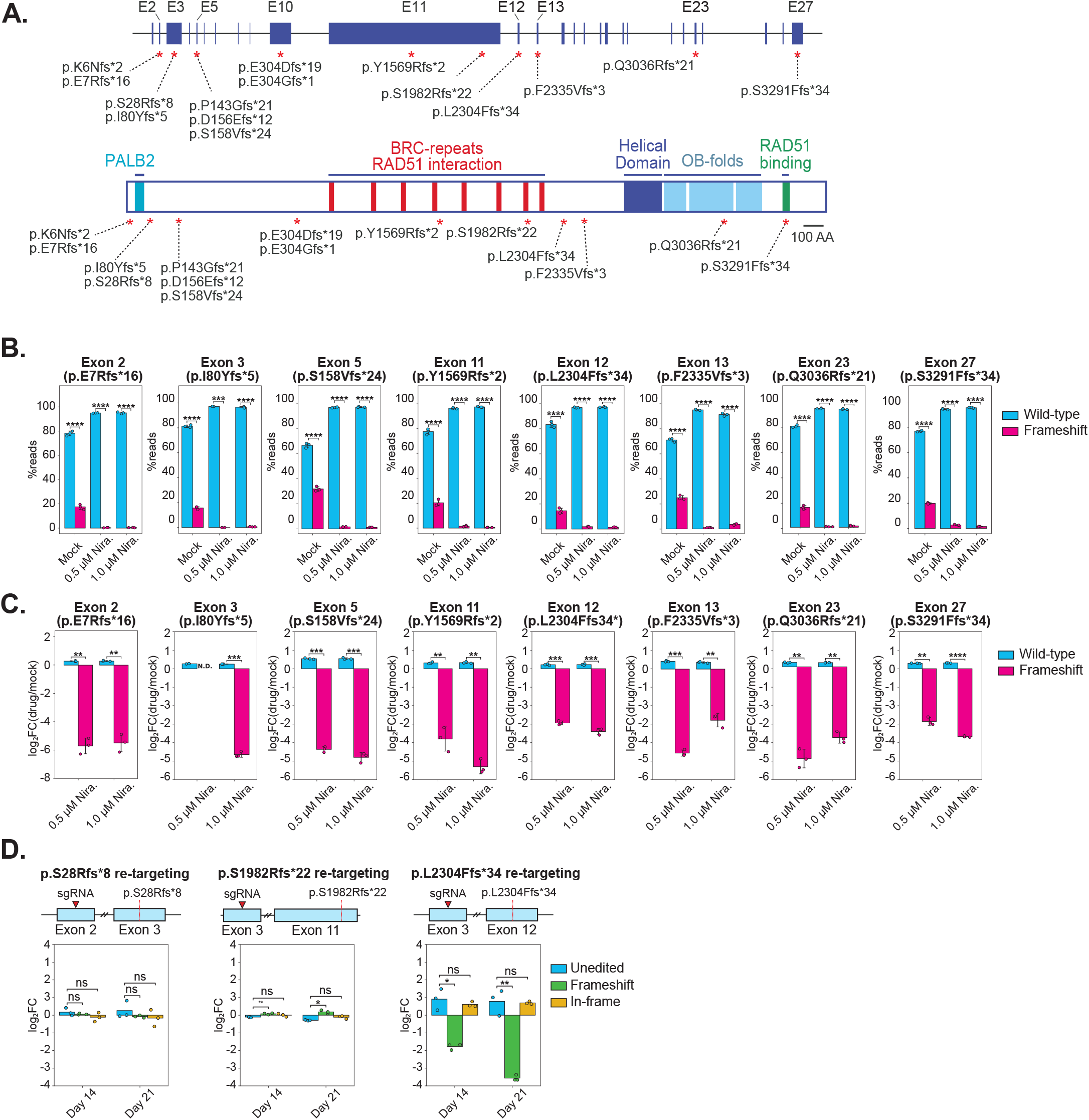
Engineered *BRCA2* frameshift mutations confer PARP inhibitor sensitivity and reveal partially functional isoforms. (A) Schematic representation of the *BRCA2* gene (top) and protein (bottom). The positions of engineered frameshift mutations are indicated by red asterisks. Key functional domains are annotated on the protein, including the PALB2 interaction domain (teal), BRC repeats/RAD51 interaction region (red), helical domain (dark blue), OB-folds (light blue), and the C-terminal RAD51 interaction domain (green). (B–C) Clonal *BRCA2* frameshift cell lines (pink) were mixed with wild-type *BRCA2* loHAP cells (blue) at a 50:50 ratio and treated with mock media, 0.5 μM niraparib, or 1 μM niraparib for 5 days (n = three technical replicates). Allele frequencies were quantified by NGS of the region surrounding the primary frameshift mutation. (B) Mean frequencies following treatment with mock, 0.5 μM, or 1 μM niraparib media across three replicates. (C) Average log2 fold change values for each allele type corresponding to the experiment shown in panel B. (D) Retargeting experiment in *BRCA2* frameshift clones. Each frameshift cell line was re-edited with an sgRNA targeting either exon 2 or 3. Schematics above each plot indicate the location of the primary frameshift mutation (red dashed line) and the sgRNA used for re-targeting (red triangle). Average log2 fold change values were calculated from the frequencies of wild-type (blue), frameshift (green), and in-frame (yellow) alleles on days 14 and 21, relative to day 7 (n = 3 biological replicates). Statistical significance was assessed using an unpaired two-sided Welch’s t-test comparing replicate mean %reads or log2 fold change between wild-type and mutant alleles within each condition. Statistical significance is indicated as follows: non-significant (NS), not detected (ND), ***P < 0.001, **P < 0.01, *P < 0.05.

To rigorously establish the p.L2304Ffs*34 (exon 12) allele as hypomorphic, this clone, along with the p.S28Rfs*8 (exon 3) and p.S1982Rfs*22 (exon 11) clones, were re-edited using an sgRNA targeting exon 2 or exon 3, which encodes the essential PALB2 interaction domain^6^. If an allele still produces a functional protein, then disrupting this domain should impair BRCA2 activity and lead to depletion in secondary frameshift alleles over time. Clones that harbor true BRCA2 LOF alleles (such as p.S28Rfs*8 and p.S1982Rfs*22) should show no additional sensitivity to re-targeting; consistently, neither clone exhibited variations in frameshift allele frequency when edited with an sgRNA targeting exons 2 or 3 (Fig. 1D). Conversely, the p.L2304Ffs*34 allele was sensitive to re-targeting in exon 3, indicating that this clone expressed a partially functional BRCA2 protein. Although this isoform retains some activity, it was insufficient to confer PARPi resistance. This is consistent with previous reports describing *BRCA2* isoforms lacking exon 12 that maintain partial BRCA2 function^26,27^.

We next asked whether the p.I80Yfs*5 clonal frameshift cell line could acquire PARPi resistance spontaneously in the absence of secondary editing. To accomplish this, three biological replicates of 5 × 10^6^ cells were treated with 1 µM niraparib for one week. Upon completion of PARPi treatment, no surviving colonies were observed. Together, these results demonstrate that frameshift mutations engineered across *BRCA2* are either null or hypomorphic and confer niraparib sensitivity, establishing a broadly representative isogenic platform for profiling PARPi-resistant reversion mutations.

### Local DNA sequence constrains the formation of *BRCA2* restoration mutations

Reversion mutations do not arise randomly; they occur predominantly within 100 bp upstream or downstream of the primary frameshift mutation and, when deletion-mediated, typically contain 2– 6 bp of microhomology at the breakpoint^17,28^. This pattern suggests that both the position of the initiating DNA lesion and the local sequence context influence the spectrum of secondary mutations that can restore the *BRCA2* reading frame. This observation is consistent with broader analyses of Cas9-induced DNA break repair, which show that repair outcomes are often biased by local sequence context and microhomology rather than being randomly distributed^29,30^, supporting the use of Cas9-induced DSBs to model how local sequence features shape *BRCA2* restoration events.

To define how local DNA constraints influence *BRCA2* restoration events, we first edited the PARPi-sensitive p.I80Yfs*5 allele in exon 3 using four sgRNAs positioned both upstream and downstream of the primary frameshift mutation (−113, −56, −41, and +33 bp; Fig. 2A). All four sgRNAs generated genetic restoration alleles and their frequency relative to frameshift alleles increased upon niraparib treatment, indicating enrichment of PARPi-resistant reversion alleles (Fig. 2B). A total of 395 unique genetic restoration alleles were detected at day 7, indicating that our system can assay these events at scale and increasing the likelihood that clinically relevant reversions will be captured (Fig. 2C, D). The frequency and spectrum of these alleles also differed markedly by sgRNA cut site: sgRNA #23, which cuts closest to p.I80Yfs*5, produced the greatest frequency and diversity of genetic restoration alleles, whereas sgRNA #10, which generated a DSB more than 100 bp upstream, yielded fewer unique restoration alleles (Fig. 2B, C).

**Figure 2:**
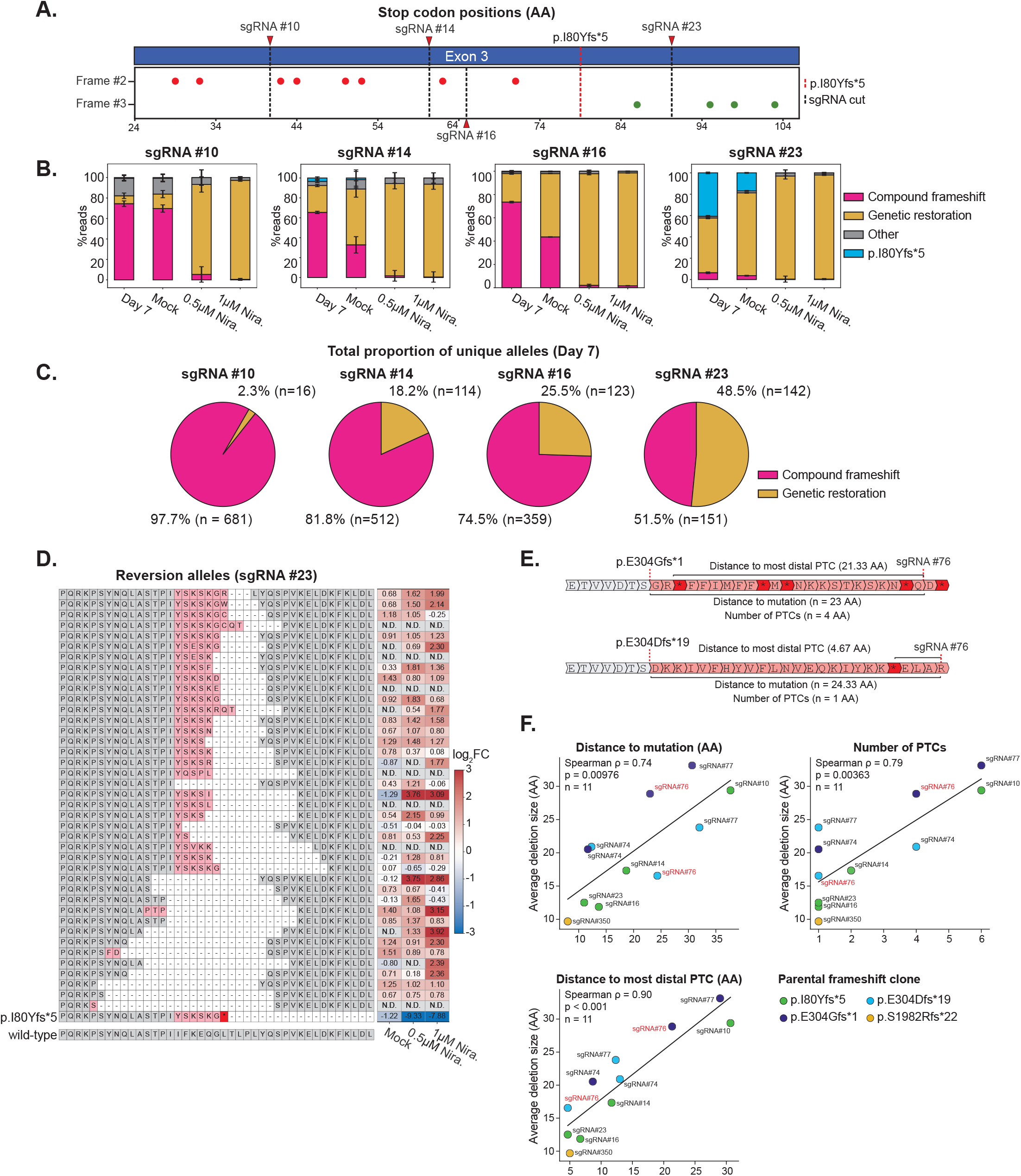
Local DNA sequence dictates BRCA2 restoration frequency and spectrum. (A) Schematic of the editing strategy used in this experiment. Four sgRNAs targeting sites upstream or downstream of the primary p.I80Yfs*5 frameshift mutation (−113, −56, −41, and +33 bp) were delivered by individual nucleofections. The positions of these sgRNAs and the p.I80Yfs*5 allele are shown relative to PTCs in frame 2 (green) and frame 3 (red). (B) Average allele frequencies for each sgRNA on day 7, and under mock, 0.5 μM niraparib, and 1 μM niraparib treatment for three biological replicates. Four allele classes were observed: edited alleles that did not restore the reading frame (compound frameshifts, pink), edited alleles that restored the reading frame (genetic restoration, yellow), unmodified p.I80Yfs*5 alleles (blue), or alleles that could not be computationally resolved (grey). Stacked bars show mean values, and error bars indicate standard deviation across three biological replicates. (C) Pie charts showing the average percentage of unique alleles across all three biological replicates on day 7, with compound frameshift mutations shown in pink and restoration alleles shown in yellow. (D) The left panel shows a heatmap of translated PARPi-resistant reversion alleles that were present in the 1uM niraparib condition at a mean frequency of 0.05% or higher, the unmodified p.I80Yfs*5 frameshift allele, and a wild-type translation for reference. The right panel shows the average log2 fold change in allele frequency relative to day 7 across three biological replicates. Alleles not detected (N.D.) in a particular condition or at day 7 are indicated in gray. (E) A representative schematic of the quantified sequence parameters shown in F for two exon 10 frameshift clones relative to the sgRNA#76 cut position (p.E304Gfs*1, top; p.E304Dfs*19, bottom). All distance parameters were measured at the nucleotide level and converted to amino acid equivalents for visualization and analysis. (F) Average restoration deletion size in the 1 μM niraparib-treated condition was compared with the distance between the parental frameshift mutation and sgRNA cut site (top left), the number of intervening premature termination codons (PTCs; top right), and the distance from the sgRNA cut site to the most distal intervening PTC (bottom left). Measurements include data from the p.I80Yfs5 parental clone (exon 3) as well as additional exon 10 frameshift clones, p.E304Gfs*1 and p.E304Dfs*19, generated in alternative reading frames, and the exon 11 p.S1982Rfs*22 frameshift clone. Points are colored according to the parental frameshift clone, as indicated. sgRNA#76 is colored in red for reference. Associations were assessed using Spearman rank correlation; Spearman’s ρ and two-sided *P* values are shown. Linear trend lines are included for visualization only. Each editing experiment was conducted with three biological replicates.

Two features likely underlie these differences in the frequency and spectrum of genetic restoration alleles: the distance between the primary mutation and the secondary DNA lesion, and the number and distribution of premature termination codons (PTCs) in the intervening sequence, which must be removed to restore the protein reading frame. To determine whether these two features influenced restorations at other loci in *BRCA2*, our analysis was extended to include three more independent genotypes, including a clone carrying the clinically relevant p.S1982Rfs*22 allele, and two clones carrying the p.E304Gfs*1 or p.E304Dfs*19 alleles, which occur at the same position in exon 10 but shift translation into alternative reading frames (Fig. 2E). Because these alleles produce distinct neopeptide tails with varying numbers of PTCs and can be edited at the same distance from the primary mutation, they allowed us to distinguish the contribution of PTC architecture from that of physical distance. The average size of PARPi-resistant reversion deletions generally scaled with both the number of intervening PTCs and the average distance between the primary frameshift mutation and the sgRNA cut site (Fig. 2F). Likewise, the reversion deletion size was most strongly correlated with the distance from the secondary DNA lesion to the most distal intervening PTC before the primary mutation (Spearman ρ = 0.90; Fig. 2F). This relationship was particularly evident in the exon 10 frameshift mutants: where sgRNA #76 produced substantially larger average reversion deletions in the p.E304Gfs*1 background than in the p.E304Dfs*19 background despite an equivalent sgRNA-to-frameshift distance. This difference likely reflects variations in the number of intervening PTCs in the two reading frames, with four PTCs in p.E304Gfs*1 and one PTC in p.E304Dfs*19. Together, these findings indicate that the frame-specific PTC landscape, along with the location of the secondary DNA lesion, constrains the size of BRCA2 reversions recovered after PARPi selection, although additional features such as cutting efficiency, microhomology availability, and local DNA repair preferences may also shape reversion frequency.

### Reversion mutations are unevenly distributed across the BRCA2 protein

Local sequence features alone do not explain the variations in reversion frequency observed across *BRCA2*^17^, and biological interpretation of this positional bias is limited by the scarcity of sequenced PARPi-resistant cases and by variations in genetic background, treatment regimens, and cancer types. To isolate the biological principle underlying these variations, we introduced designer genetic restoration mutations both upstream of the previously defined hotspot (exon 5) and within the cold spot (exons 17 and 20; DNA binding domain), and assayed their ability to confer PARPi resistance in a pooled setting^17^ (Fig. 3A). These genetic restorations were engineered through a dual-editing CRISPR-Cas9 strategy, in which frameshift alleles were introduced at an initial position within an exon using the first sgRNA. Cells were then cultured for one week and edited again with a second sgRNA, proximal to the first, to restore the reading frame in a subset of the alleles (Fig. 3B). Edited cell populations were then expanded for two weeks before being treated with either mock or 1 µM niraparib media. Assessing these genetic restoration mutations using this bulk-editing strategy allows direct comparison of their fitness relative to frameshift and wild-type alleles. To generate genetic restorations in both reading frames, oligo pools containing designer 1- or 2-bp frameshift alleles were delivered as repair templates with the first sgRNA and subsequently re-edited with the second sgRNA to restore BRCA2 expression (Fig. 3B). Given our previous result that the frequency of PARPi-resistant reversions is inversely correlated with the number of intervening PTCs, we introduced synonymous mutations in all PTCs located between the first and second sgRNA cut sites (Fig. 3C). This design increased the recovery of engineered genetic restoration alleles and enabled direct comparison of their fitness relative to frameshift and wild-type alleles following treatment with mock or 1 μM niraparib.

**Figure 3:**
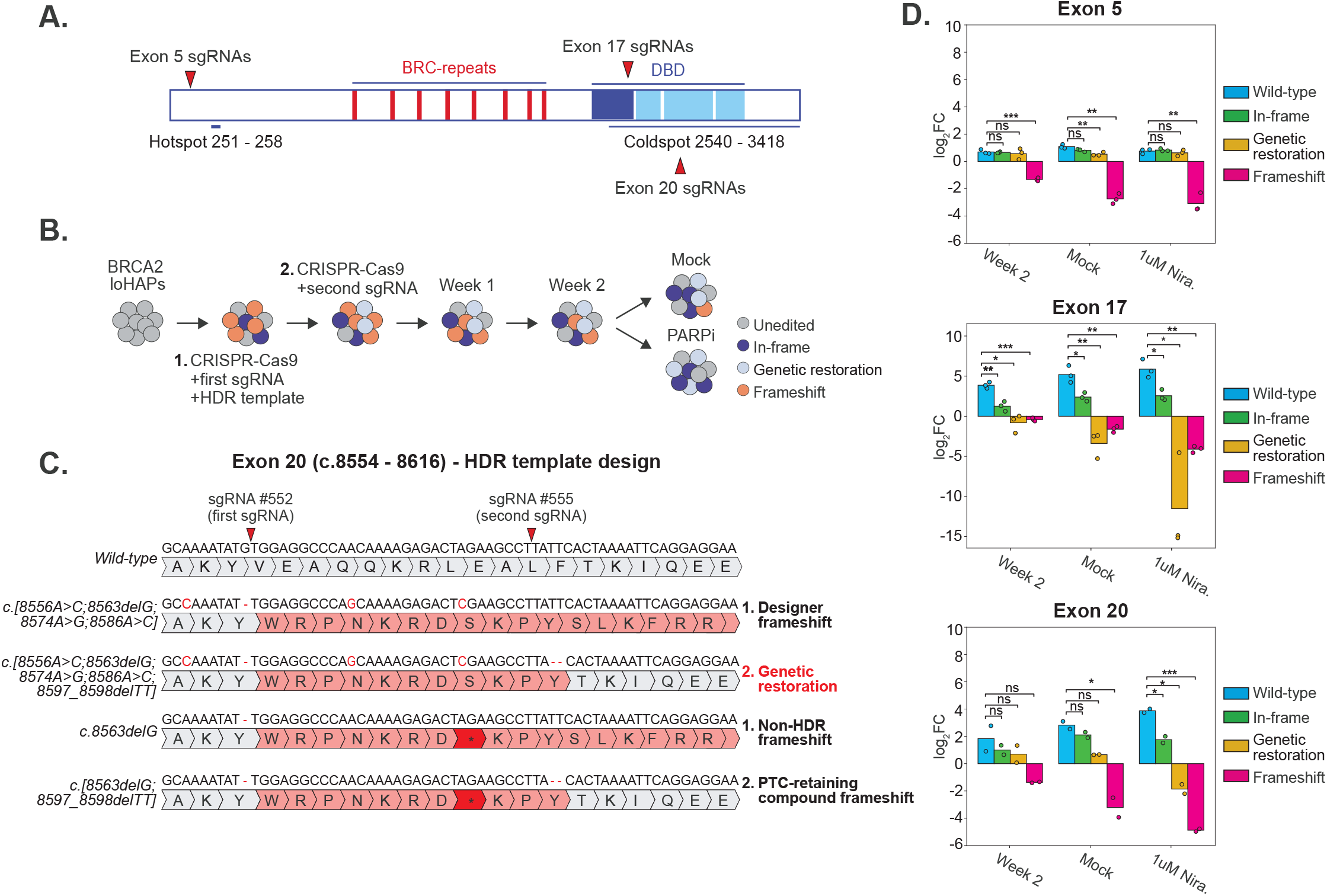
PARPi-resistant *BRCA2* reversions exhibit a positional bias. (A) Schematic of the BRCA2 protein highlighting the N-terminal reversion hotspot (amino acids 251–258) and the C-terminal cold spot (amino acids 2540–3418), along with the positions of BRC repeats and the DNA-binding domain (DBD). sgRNAs targeting exon 5 (outside the cold spot) and exons 17 and 20 (within the cold spot) are indicated. (B) Experimental design for generating restoration mutations using a dual CRISPR–Cas9 editing strategy. In Step 1, designer frameshift alleles were introduced by nucleofection with the first sgRNA and an HDR template containing the intended frameshift and synonymous substitutions that remove intervening PTCs. Cells were expanded for one week before nucleofection with a second sgRNA in Step 2 to generate restoration alleles. Three biological replicates were then expanded for two weeks and treated with mock or niraparib media.. (C) Representative exon 20 schematic illustrating editing outcomes from Steps 1 and 2 in (B). HDR-mediated editing in Step 1 generates a designer frameshift allele in which intervening PTCs are removed. Editing with the second sgRNA in Step 2 can then restore the *BRCA2* reading frame, generating a genetic restoration allele. In contrast, non-HDR alleles retain an intervening PTC and can yield a PTC-retaining compound frameshift after secondary editing. These outcomes illustrate the rationale for removing intervening PTCs from the designer HDR templates. (D) Average log2 fold change in allele frequencies for wild-type (blue), in-frame (green), genetic restoration (yellow), and frameshift (pink) alleles in exons 5, 17, and 20 at week 2, under mock, or under 1 μM niraparib treatment relative to day 7. Restoration alleles are identified based on a unique neopeptide generated by a double-editing event. In-frame alleles are those that restore the reading frame but do not contain a distinct neopeptide, including single in-frame editing events, dual in-frame events generated by both sgRNAs, and large in-frame editing events that encompass both sgRNA cut sites. Frameshift alleles are defined as those with either a single or double edit that do not restore the reading frame. Three biological replicates were analyzed for exons 5 and 17. The exon 20 experiment was initiated with three biological replicates, but one replicate was excluded because the day 7 sample failed sequencing-depth quality control and therefore could not support reliable log2 fold-change calculation. Statistical significance was assessed using an unpaired two-sided Welch’s t-test comparing replicate-level log2 fold-change values between wild-type and mutant allele classes within each condition. Because only two biological replicates were available for exon 20, these results are interpreted as supportive rather than definitive. A pseudo count equal to one-half of the lowest nonzero allele count was added before calculating log2 fold-change values. Statistical significance is indicated as follows: non-significant (NS), not detected (ND), ***P < 0.001, **P < 0.01, *P < 0.05.

Frameshift alleles in exon 5 were depleted following 1 µM niraparib treatment, whereas genetic restoration alleles were enriched, indicating that these events constituted true PARPi-resistant reversion mutations. By contrast, both genetic restoration and frameshift alleles introduced in exons 17 and 20 within the cold spot were strongly depleted after niraparib treatment (Fig. 3D). These results suggest that genetic restoration events are subject to an additional layer of functional selection, such that only those that preserve essential BRCA2 functions, including those encoded within the C-terminal DNA-binding domain, retain sufficient activity to behave as PARPi-resistant reversions. Because these regional variations in *BRCA2* reversion frequency were recapitulated in our isogenic system, they likely reflect an intrinsic positional bias that can be distinguished from differences in genetic background, treatment history, or cancer type.

### BRCA2’s PALB2 interaction domain is required for PARPi resistance

To directly test whether disruption of a defined functional domain could limit which genetic restoration events confer PARPi resistance, we systematically evaluated genetic restoration events in exon 3, which contains the C-terminal half of the PALB2 interaction domain^6^. Mutation or deletion of amino acids (AA) 10–40, which encode this domain, impairs or ablates BRCA-mediated HR, and renders cells PARPi sensitive^6,21^. However, PARPi-resistant reversions were readily generated elsewhere in this exon, as demonstrated in Fig. 2. To systematically map genetic restoration events in this region, we designed a lentiviral library containing sgRNAs targeting all NGG PAM sites (n = 19) in exon 3, which was transduced at low multiplicity of infection into the PARPi-sensitive p.I80Yfs*5 cell line (exon 3). This cell line harbors a frameshift mutation 39 residues downstream of the PALB2 interaction domain. Following one week of puromycin selection to select for cells carrying the sgRNA construct, cells were nucleofected with Cas9 protein and expanded for one week (Fig. 4A). Cells were then treated with either mock or 1 µM niraparib for an additional week to enrich PARPi-resistant reversion alleles. Samples were collected at week 1 from the mock- and drug-treated conditions and analyzed by NGS to quantify sgRNA and allele frequencies at the exon 3 locus.

**Figure 4:**
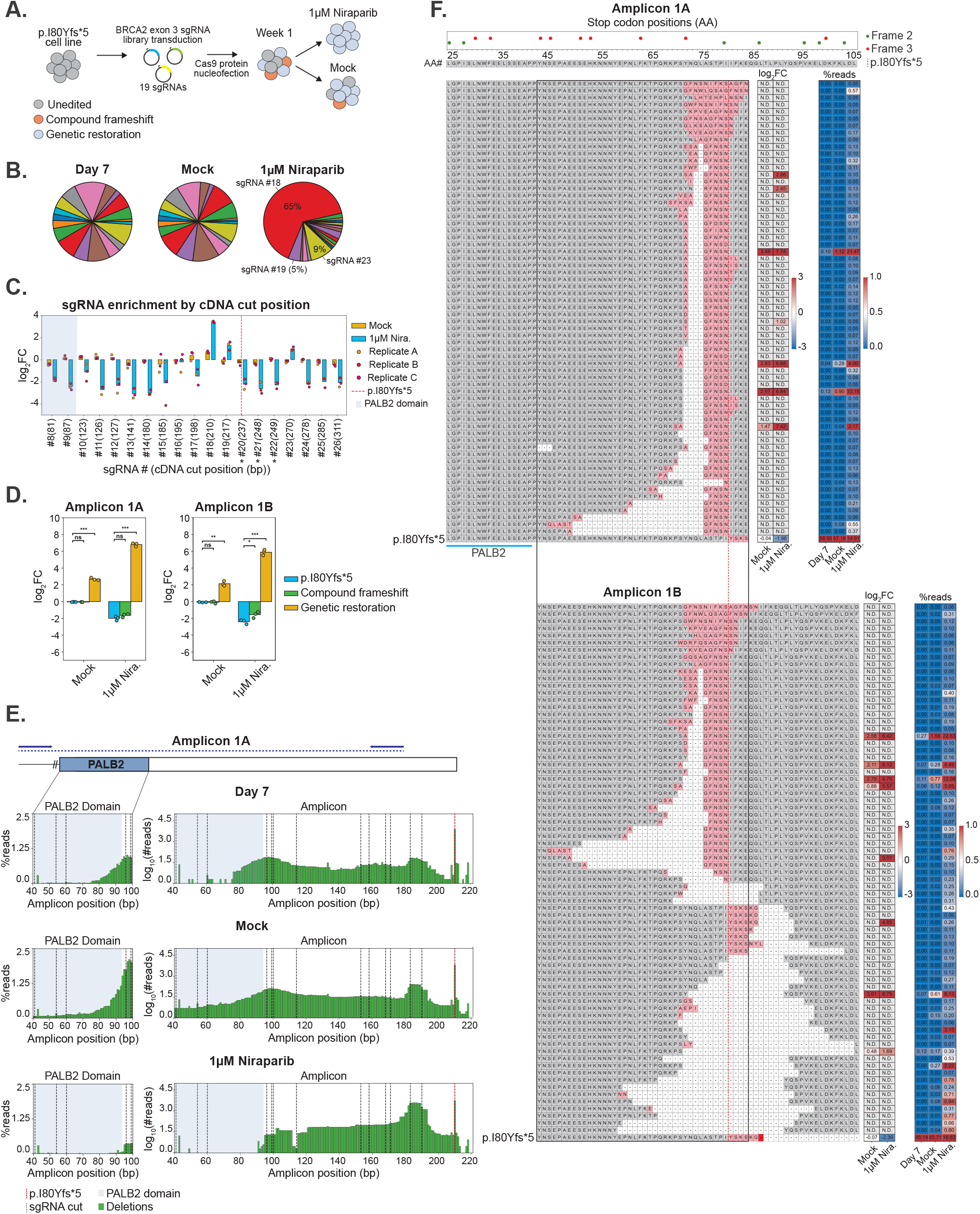
The PALB2 interaction domain is required to confer PARPi resistance. (A) Schematic of the pooled CRISPR–Cas9 restoration tiling strategy. A lentiviral sgRNA library containing 19 sgRNAs targeting *BRCA2* exon 3 was transduced into the PARPi-sensitive p.I80Yfs*5 cell line, followed by Cas9 protein nucleofection, one week of expansion, and treatment with either mock or 1 μM niraparib. Genetic restoration alleles (light blue) were enriched relative to unedited (gray) and compound frameshift (orange) alleles under both conditions, with stronger enrichment under niraparib selection. (B) Pie charts showing average sgRNA representation at day 7 and after mock or 1 μM niraparib treatment. sgRNAs #18 and #23 comprised the majority of the population after PARPi treatment. (C) Average log2 fold change in sgRNA abundance, plotted by cDNA cut position, for mock (yellow) and 1 μM niraparib (blue) conditions relative to day 7. sgRNAs #20–22 are italicized and marked with an asterisk to indicate that cutting is blocked by the parental frameshift mutation. Individual biological replicates are shown as dots (n = 3). A red dashed line indicates the position of the p.I80Yfs*5 mutation, and the PALB2 interaction domain is shaded in light purple. (D) Average log2 fold change of allele classes recovered from amplicons 1A and 1B under mock or 1 μM niraparib treatment relative to day 7. The p.I80Yfs*5 allele is shown in blue, compound frameshifts in green, and genetic restoration alleles in yellow. Statistical significance was assessed using an unpaired two-sided Welch’s t-test comparing the average replicate log2 fold change between p.I80Yfs*5 and mutant allele types within each condition (n = 3 biological replicates). Statistical significance is indicated as follows: non-significant (NS), not detected (ND), ***P < 0.001, **P < 0.01, *P < 0.05. (E) Average deletion frequency per base pair across amplicon 1A at day 7 and after mock or niraparib treatment. Left panels show the unnormalized average deletion frequency within the PALB2 interaction domain. The right panels show the log10-fold change in the normalized number of deletion reads to better visualize editing efficiency on day 7. A red dashed line indicates the position of the p.I80Yfs*5 allele, sgRNA cut sites are indicated by gray dashed lines; the PALB2 interaction domain is indicated by light purple shading; and deletions are shown in green. (F) Heat maps showing the spectrum and frequency of PARPi-resistant reversion alleles detected in amplicons 1A and 1B. Stop codon positions in reading frames 2 and 3 are indicated above the heat maps. The positions of the PALB2 interaction domain and the p.I80Yfs*5 mutation are marked. Only restoration alleles with an average frequency greater than 0.05% in the 1 μM niraparib condition are shown. The left panel displays representative translations with neopeptides highlighted in red. The middle panel shows the average log2 fold change in allele frequency under mock and 1 μM niraparib relative to day 7. The right panel shows average allele frequencies at day 7 and after mock or niraparib treatment.

Consistent with the positional bias described above, sgRNAs that cut closest to the p.I80Yfs*5 mutation (sgRNAs #18, #19, #23) were enriched under 1 µM niraparib treatment (Fig. 4B, C). In contrast, sgRNAs targeting the PALB2 interaction domain (cDNA positions 68–120) were not enriched (Fig. 4C). To determine whether editing events were generated within the PALB2 interaction domain despite the lack of sgRNA enrichment after PARPi treatment, the exon 3 locus was sequenced using amplicons spanning regions upstream and downstream of the p.I80Yfs*5 mutation (amplicons 1A and 1B, respectively). NGS analysis revealed that PARPi-resistant reversion mutations in both amplicons were enriched in the niraparib-treated population relative to the unmodified p.I80Yfs*5 allele and compound frameshift alleles that did not restore the reading frame (p < 0.001, Fig. 4D).

Unlike concurrent clinical studies^31,32^, which typically identify *BRCA2* reversion mutations after drug resistance has emerged, our platform detects restoration alleles before selection and can determine which are subsequently enriched under PARPi treatment. This approach provides evidence that a specific reversion mutation contributes to PARPi resistance, rather than merely behaving as a passenger mutation. To confirm that editing events were introduced in the PALB2 interaction domain, the average deletion frequency per nucleotide was calculated for amplicon 1A on day 7 (Fig. 4E, left panels). Nucleotides 42–92 in amplicon 1A, which encompass the PALB2 interaction domain, exhibited a cumulative per-position deletion frequency of approximately 7%, with editing events concentrated at the 3′ end of the domain. This finding confirmed that edits were generated in this region, a subset of which could theoretically produce restoration events that remove part or all of the PALB2 interaction domain. However, the average deletion frequency within the PALB2 interaction domain decreased significantly following treatment with 1 µM niraparib. Frequencies in the remainder of the amplicon either increased or remained unchanged (Fig. 4E, right panels). Similarly, PARPi-resistant reversion alleles with an average allele frequency greater than 0.05% in the 1 µM niraparib condition did not extend into the PALB2 interaction domain (AA 24–40) (Fig. 4F), with the largest enriched reversion beginning three AA downstream of the domain. Reversions outside of this region were highly diverse and spanned both upstream and downstream of the p.I80Yfs*5 mutation (Fig. 4F). Thus, while mutations were readily introduced across exon 3, genetic restoration alleles extending into the PALB2 interaction domain were not observed following PARPi selection. These findings demonstrate that the PALB2 interaction domain is required to confer PARPi resistance and provide an example of how local functional domain architecture can shape regional reversion patterns.

### Interexonic restoration events adhere to shared principles

Interexonic restoration events re-establish an in-frame *BRCA2* transcript across exon boundaries through either genetic restoration of the reading frame or altered splicing. These events are likely under-sampled and misclassified in clinical datasets, such limitations may arise in part from the technical constraints of clinical sequencing, as restricted amplicon size can exclude distant secondary mutations and genomic sequencing alone cannot determine whether a secondary mutation produces a functional *BRCA2* reversion^33^. To test whether interexonic restoration events could be generated and positively selected in our *BRCA2* loHAP system, we re-edited the hypomorphic p.L2304Ffs*34 (exon 12) clone with an sgRNA targeting the exon 13 splice donor, as clinical PARPi-resistant reversions have been reported at the exon 13/14 junction^18^. Following nucleofection with this exon 13 sgRNA and a one-week expansion, cells were treated with either mock or 1 µM niraparib. cDNA was synthesized from each condition, and the region spanning exons 12–14 was amplified alongside genomic DNA surrounding exon 13. NGS of the cDNA revealed that interexonic reversions were significantly enriched relative to compound frameshift mutations in the 1 µM niraparib-treated condition (p < 0.01, Supplemental Fig. 1A). Analysis of translated alleles identified two mechanisms of restoration (Supplemental Fig. 1B). One class restored the reading frame across exons 12-14 at the DNA level, while the second arose through exon 13 skipping. DNA-level analysis of PARPi-enriched alleles further supported splicing-mediated rescue at this locus, as several enriched alleles deleted or modified the canonical splice donor (Supplemental Fig. 1C). These results show that both genetic and splicing-mediated restoration mechanisms can generate interexonic *BRCA2* reversions, and that exon 13 is dispensable for PARPi resistance in BRCA2-deficient cells.

We next asked whether the sequence-based constraints could explain where interexonic restoration events arise. To address this, we generated a lentiviral sgRNA library targeting every NGG PAM within *BRCA2* coding exons 2-10. This region was selected because it lies outside the reversion cold spot^17^ and is prone to alternative splicing. This N-terminal domain (NTD) library was transduced into three exon 5 frameshift clones with two distinct local sequence architectures. The p.S158Vfs*24 and p.P143Gfs*21 alleles generated neopeptide tails in frame 2 that extended into exon 7, whereas the p.D156Efs*12 allele generated a shorter neopeptide tail in frame 3 that terminated in exon 6. We therefore predicted that restoration of the first two alleles could occur through more distal events extending into exon 7, whereas restoration of p.D156Efs*12 would be more frequent in exon 6. To test this, transduced and selected cell populations were nucleofected with Cas9 protein and then treated with either mock or 1 µM niraparib. Analysis of cDNA spanning exons 4 - 8 revealed genetic interexonic restoration events whose distribution differed according to the reading frame of the parental frameshift (Fig. 5A - C). Genetic restorations in the p.P143Gfs*21 and p.S158Vfs*24 (frame 2) clones occurred more frequently in downstream exons 6 and 7, whereas those in the p.D156Efs*12 (frame 3) clone remained confined to exon 6. Consistent with this observation, NGS analysis of sgRNA frequency in the PARPi-treated edited cell populations revealed sgRNAs targeting exon 7 were the most enriched in the p.P143Gfs*21 and p.S158Vfs*24 clones in reading frame 2, while sgRNAs targeting exon 6 were more enriched in the p.D156Efs*12 clone in reading frame 3 (Fig. 5D). Together, this pattern suggests that interexonic restoration is shaped by the same sequence-based constraints as local intraexonic restoration, with the parental mutation’s reading frame and local PTC density determining neopeptide tail length and thereby the range over which downstream restoration can occur.

**Figure 5:**
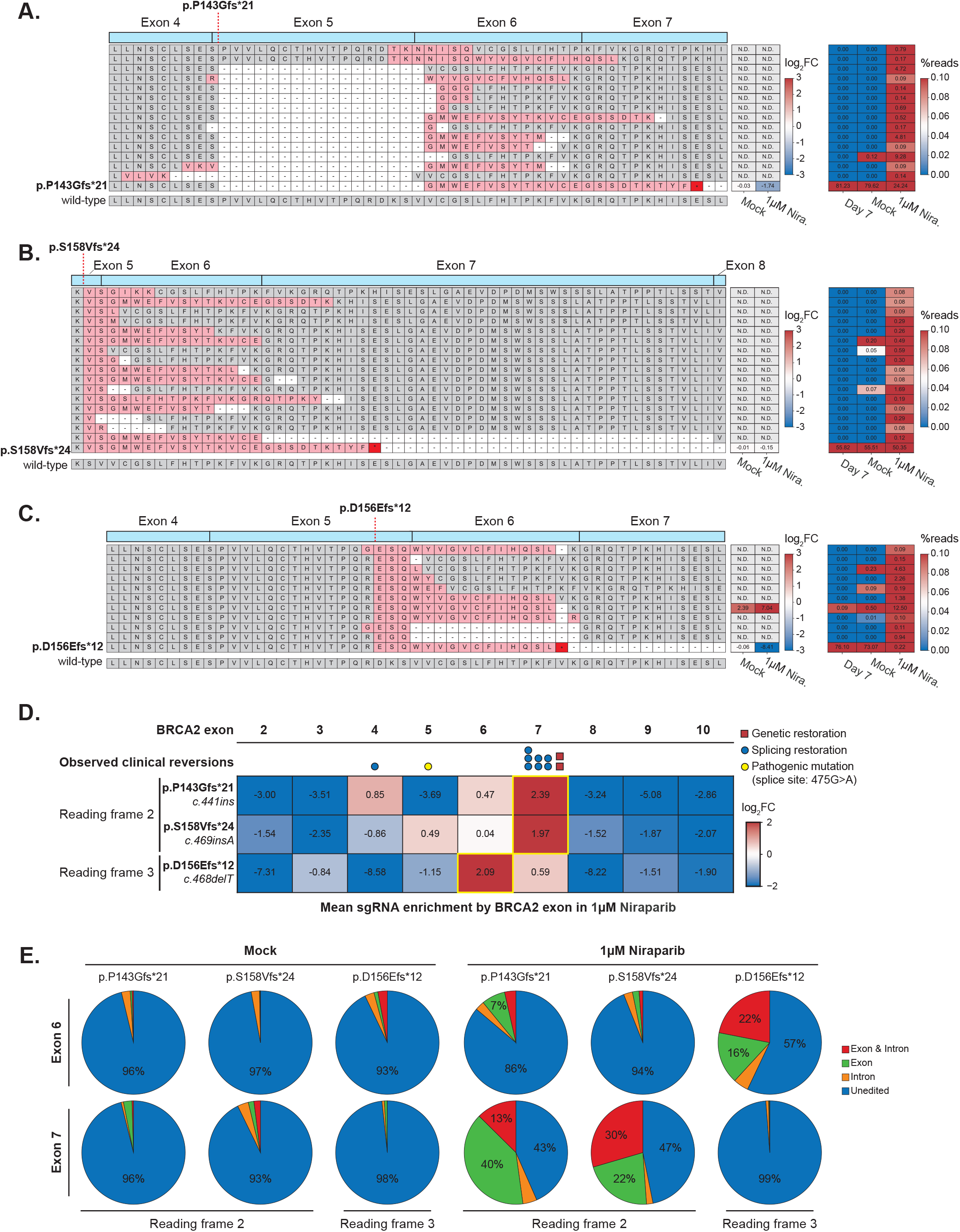
Interexonic *BRCA2* restorations are shaped by reading frame and local splicing architecture and arise through distinct mechanisms. Allele frequency analysis from the BRCA2 N-terminal domain (NTD) lentivirus editing experiment targeting *BRCA2* exons 2 - 10. (A - C) PARPi-resistant reversion allele enrichment in (A) p.P143Gfs*21, (B) p.S158Vfs*24, and (C) p.D156Efs*12 edited parental backgrounds following niraparib selection at the cDNA level. Translated sequences are aligned across the indicated *BRCA2* exons, with the position of the parental frameshift mutation indicated by the red dashed line. Neopeptide sequences are highlighted in red. For each allele, the adjacent heatmap shows the mean log2 fold change in allele frequency under mock or 1 μM niraparib treatment relative to day 7, and the rightmost heatmap shows the mean percentage of sequencing reads at day 7 and following mock or niraparib treatment. Alleles not detected in a given condition are indicated as N.D. The parental frameshift and wild-type sequences are shown for reference. (D) Mean log2 fold change in sgRNA abundance following 1 μM niraparib treatment relative to mock, grouped according to the *BRCA2* exon containing the sgRNA cut site. The experiment was performed in three parental *BRCA2* frameshift backgrounds: p.P143Gfs*21 and p.S158Vfs*24, which have a neopeptide tail in reading frame 2, and p.D156Efs*12, which has a neopeptide tail in reading frame 3. The HGVS cDNA names are shown below the parental frameshift names for reference. Values represent the mean enrichment of sgRNAs targeting each exon across three biological replicates. Yellow outlines indicate the most strongly enriched exon for each parental frameshift background. The p.P143Gfs*21 and p.S158Vfs*24 backgrounds preferentially enrich sgRNAs targeting exon 7, whereas the p.D156Efs*12 background preferentially enriches sgRNAs targeting exon 6. Above the heatmap, previously observed drug-resistant clinical *BRCA2* reversions are plotted according to exon. Reversions arising through genetic restoration of the reading frame across exons are shown as red squares, and those arising through splicing-mediated restoration as blue circles. The pathogenic *BRCA2* splice-site variant *c.475G>A* is represented by a yellow circle. (E) Pie charts showing the mean proportion of alleles within the genomic regions spanning exon 6 (top) or exon 7 (bottom), including the entire exon and 50 bp of flanking intronic sequence on either side. Alleles were classified as unmodified or containing indels (insertions and deletions) occurring exclusively within the exon, exclusively within the intronic regions, or spanning both the intron and exon. Substitutions were excluded from this analysis.

Interestingly, the local transcript architecture in each parental frameshift clone influenced both the frequency and pattern of splicing-mediated interexonic restorations and their propensity to undergo genetic restorations in exon 4. First, the p.P143Gfs*21 clone, which contains a 14 bp insertion in the canonical splice donor of exon 5, causing out-of-frame skipping of this exon, was the only clone with genetic restorations in exon 4 (Fig. 5A). The absence of exon 5 from the parental transcript likely explains why sgRNAs targeting exon 5 were not enriched under PARPi treatment (Fig. 5D). Notably, sgRNAs were enriched in the same exons in which reversions occurred in a patient carrying the exon 5 splice-site mutation *c.475G>A*^31^, suggesting concordance between the regions susceptible to reversion in our experimental system and in a clinical setting. When exon 5 was retained, as in the p.S158Vfs*24 clone, splicing-mediated restorations resulted in skipping of exon 7 (Fig. 5B). Finally, only the p.D156Efs*12 clone had splicing-mediated restorations that resulted in the skipping of exon 6 (Fig. 5C). To determine whether these exon-skipping events were associated with changes at the corresponding splice junctions, we sequenced the genomic regions surrounding exons 6 and 7. NGS of the PARPi-treated edited cell populations revealed that indels overlapping the canonical intron-exon boundaries in exon 7 were enriched in the p.S158Vfs*24 clone, while those overlapping with exon 6 were enriched in the p.D156Efs*12 clone (Fig. 5E). Deletion or mutation of a splice donor or acceptor in these regions could disrupt splice-site recognition and provide a mechanism for the exon-skipping events observed in these clones. Taken together, these findings indicate that the reading frame of the parental transcript determines which downstream exon-skipping event restores an in-frame *BRCA2* transcript, explaining the distinct exon 7- and exon 6-skipping patterns observed in the p.S158Vfs*24 (frame 2) and p.D156Efs*12 (frame 3) backgrounds, respectively.

Next, we asked whether interexonic restoration events are subject to the same secondary functional selection that determines which events confer PARPi resistance, and can therefore be classified as interexonic reversions. Single-guide editing experiments were performed in the p.E7Rfs*16 (exon 2) and p.Q3036Rfs*21 (exon 23) clones at the exon 2/3 and exon 23/24 junctions, which overlap the PALB2 interaction domain and OB folds, respectively (Supplemental Fig. 2A, B). Although we detected interexonic restoration mutations in both settings, no viable colonies emerged under PARPi treatment. This lack of outgrowth indicates that the BRCA2 domain architecture imposes a second layer of functional selection, as observed with intraexonic restorations, such that only a subset of interexonic restoration events confer PARPi resistance and constitute reversion alleles.

### *BRCA2* exon 11 is uniquely permissive to large PARPi-resistant reversions

Reversions arising from large deletions in exon 11 are frequently observed in clinical datasets^18,28^. Interpretation of these events is complicated by discrepancies between clinical and experimental systems, in which reported exon 11 restoration events range from small local mutations to much larger deletions that remove up to six BRC repeats and most of the BRCA2 C-terminus^35,36^. Splicing-mediated restoration events in exon 11 have also been reported in cell lines^35,37^. These observations raise the question of whether the clinical and experimental spectra of exon 11 restoration events can be reconciled to define the mechanisms that generate PARPi-resistant reversions in this region, and whether exon 11 adheres to the sequence- and domain-based constraints that limit *BRCA2* reversion elsewhere in the gene. To answer this, we generated a gene-scale lentiviral sgRNA library targeting every NGG PAM site across the *BRCA2* coding sequence (n = 690, including 42 negative controls). The library was transduced into a *BRCA2* loHAP cell line carrying the exon 11 frameshift mutation p.Y1569Rfs*2. Following nucleofection and niraparib selection, sgRNAs targeting exon 11 became progressively enriched and accounted for 92% of the library in the 1 μM niraparib condition (Supplemental Fig. 3A), indicating that PARPi-resistant reversions in this background were largely confined to exon 11. A small subset of sgRNAs were also enriched in the 5’ end of exon 12 (Supplemental Fig. 4).

The two most frequent sgRNAs in the 1 μM niraparib condition (#339 and #354) accounted for 50% of the total population and cut 974 bp and 1,257 bp downstream of p.Y1569Rfs*2 in exon 11, respectively (Supplemental Fig. 3B, C). Additional sgRNAs were broadly enriched throughout exon 11, including within 200 bp of the p.Y1569Rfs*2 mutation and near the 3′ end of exon 11. This enrichment of distal sgRNAs was striking, given our earlier results showing that reversion frequency generally declines with increasing distance from the primary frameshift (Fig. 2). To define these events, we clonally isolated revertants from the 1 μM niraparib condition and characterized them at the DNA and *BRCA2* transcript levels (Supplemental Fig. 3D - G). All revertants arose from sgRNAs #339 or #354. At least three genotypes were identified from genomic DNA, including Δ2912;ins331 bp, Δ2248 bp, and one unresolved event (Supplemental Fig. 3E). However, intervening PTCs remained between each deletion breakpoint and the primary mutation, excluding direct DNA-level restoration of the reading frame as the basis for PARPi resistance in these clones. PCR amplification of exons 10 through 12 from *BRCA2* cDNA revealed that each revertant clone expressed multiple transcript isoforms, consistent with alternative splicing following disruption of the canonical exon 11 splice donor (Supplemental Fig. 3F). Four of these isoforms appeared to be similar in size across all genotypes. Sequencing of the largest isoform showed that the 3′ end of BRC4 functions as a cryptic splice donor to exon 12 (Supplemental Fig. 3G). Thus, despite distinct underlying deletions, all revertants expressed the same alternative isoform via a shared cryptic splice donor. The same splicing event was reported in a patient with treatment-resistant breast cancer, indicating that this event likely represents a conserved restoration mechanism underlying reversion^35,38^.

Our data identify exon 11 as a uniquely permissive region for *BRCA2* reversion, in which much of the exon 11 sequence, including the BRC repeats, appears relatively tolerant of large deletions. Notably, these reversion events remained confined to exons 11 and 12 rather than extending across the C-terminal DNA-binding domain, yielding a pattern that closely matches the clinically observed spectrum of *BRCA2* reversions^17^.

### *c.5946delT* (p.S1982Rfs*22) reveals conserved splicing-mediated restoration underlying PARPi resistance

To further test the extent to which these alternative splicing events can generate PARPi-resistant reversions, we next asked whether the use of cryptic splice donors extended to a clinically relevant mutant background. *c.5946delT* (p.S1982Rfs*22) is a founder mutation that occurs in approximately 1% of individuals of Ashkenazi Jewish ancestry^39^. Reversions arising from this mutation show a marked 3′ bias, occurring more frequently downstream of p.S1982Rfs*22 than upstream, in contrast to many other pathogenic *BRCA2* mutations^17^. Because this mutation is both clinically recurrent and associated with a characteristic reversion pattern, it provides a strong test of whether splicing-mediated restoration through cryptic donor usage is a general mechanism underlying reversions across independent *BRCA2* genotypes.

To test whether similar restoration patterns could be recapitulated in the p.S1982Rfs*22 background, we performed deep mutagenesis in a loHAP cell line engineered to carry this mutation. The mutant line was transduced with a lentiviral sgRNA library targeting every NGG PAM in exon 11 (n = 264), nucleofected with Cas9, and treated with 1 µM niraparib. Alkaline phosphatase (AP) staining revealed PARPi-resistant colonies emerged only in Cas9-nucleofected cells, not in the parental control population, indicating that reversions were successfully generated (Supplemental Fig. 5A). sgRNA analysis after niraparib selection identified three dominant enriched guides (#341, #355, and #392), together with broader enrichment of guides cutting within 300 bp of p.S1982Rfs*22 and within 100 bp of the 3′ end of exon 11 (Supplemental Fig. 5B, C). This enrichment pattern recapitulates the clinically reported 3′ bias associated with the p.S1982Rfs*22 founder mutation. Notably, sgRNAs #341, #355, and #392 cut within 42 bp downstream of the corresponding dominant guides identified in the p.Y1569Rfs*2 experiment (sgRNAs #339, #354, and #391, respectively) (Supplemental Fig. 3C and Supplemental Fig. 5C). The close proximity of enriched sgRNAs across these independent exon 11 frameshift alleles, which are themselves located 1240 bp apart, indicates that PARPi-resistant reversions in this region are constrained to reproducible sites and likely arise through shared underlying restoration mechanisms.

To link sgRNA enrichment with the underlying restoration mechanisms, we performed long-read nanopore sequencing on the niraparib-selected edited populations from the p.Y1569Rfs*2 and p.S1982Rfs*22 experiments. Two amplicons were sequenced to capture splicing-mediated restoration events impacting both the exon 11 splice acceptor and donor (Supplemental Fig. 6A, Fig. 6A). Alignment of reads to the wild-type BRCA2 cDNA reference revealed a broad spectrum of restoration events in both genotypes (Fig. 6B - D). None of these events were observed in the parental frameshift clones treated with PARPi, which, as in all previous experiments, did not yield spontaneous revertants. These events fell into two classes: intraexonic genetic deletions that restored the reading frame while preserving canonical splice junctions, and splicing-mediated restoration events generated by a cryptic splice donor in exon 11 and the canonical exon 12 splice acceptor. Across both backgrounds, 29 unique genetic deletions were identified, with the largest deleting seven of the BRC repeats in the p.Y1569Rfs*2 clone. Five distinct splicing-mediated restoration events were also identified. These used predicted splice donors identified by NNSPLICE 0.9^40^, supporting their classification as true splicing events rather than deletions extending to exon 12. Of the 16 predicted cryptic splice donors in exon 11, only six were predicted to generate in-frame transcripts, five of which we identified in our dataset. Three splicing isoforms were shared between the p.Y1569Rfs*2 and p.S1982Rfs*22 backgrounds, whereas the genetic deletions were largely genotype-specific. The largest of these isoforms removed all eight BRC repeats. No splicing-mediated restoration events were detected that paired the canonical exon 10 donor with a cryptic acceptor in exon 11, even though one of the six predicted cryptic splice acceptors was expected to generate in-frame transcripts in both frameshift clones (Supplemental Fig. 6B-D). Together, these results indicate that the restoration mechanisms underlying exon 11 reversion are highly conserved, that splicing-mediated restoration is directionally biased toward cryptic donor activation, and that all eight BRC repeats may be dispensable for PARPi resistance. However, because this isoform was detected alongside additional restoration transcripts in the PARPi condition, its presence does not establish whether a complete loss of BRC repeats was itself sufficient to confer PARPi resistance.

**Figure 6:**
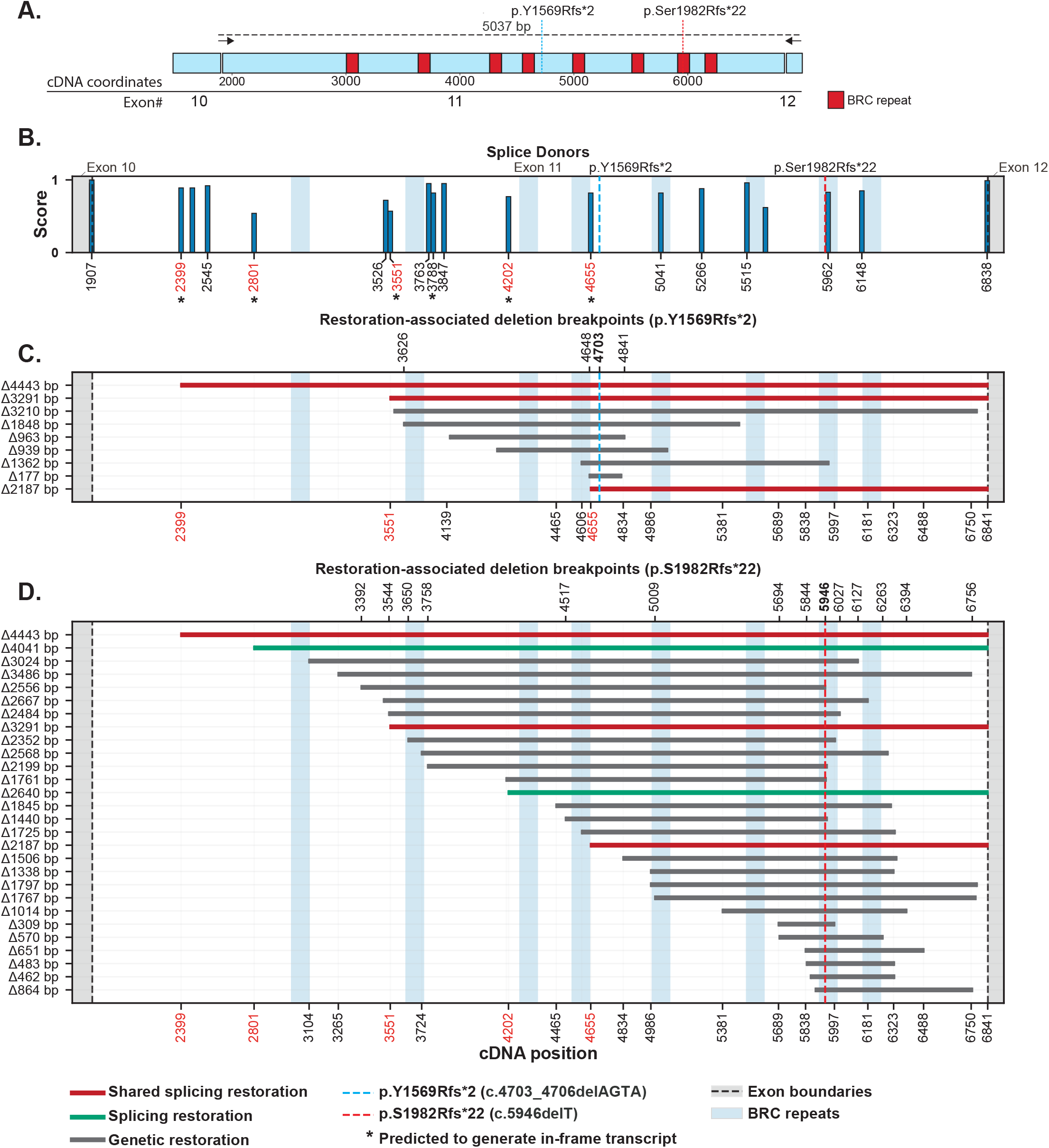
Long-read sequencing reveals conserved splice donors in exon 11. (A) Schematic of the PCR strategy used for nanopore sequencing of *BRCA2* cDNA. PCR primers were anchored at the exon 10–11 junction and in exon 12. The expected PCR product size for the wild-type transcript was 5037 bp. (B) Predicted splice donor strength across exons 10–12, plotted as a function of cDNA position. Multiple high-scoring cryptic splice donor sites were identified within exon 11. Splice-site strength predictions were generated using NNSPLICE 0.9. Alternative splice donors that generate an in-frame product are indicated by an asterisk. (C, D) Deletion breakpoints of restoration alleles identified by long-read nanopore sequencing of the cDNA amplicon isolated from the 1 μM niraparib-treated condition in the p.Y1569Rfs*2 (C) and p.S1982Rfs*22 (D) backgrounds. Genetic restorations are shown in gray, genotype-specific splicing-mediated restorations in green, and shared splicingmediated restorations detected in both genotypes in red. Coordinates of splicing-mediated restorations are indicated in red. Exon boundaries are indicated by gray shading and dashed lines, BRC repeats are shaded in blue, and the positions of the parental frameshift alleles are indicated. Three biological replicates were used to generate nanopore data.

### At least one BRC repeat is required for PARPi-resistant BRCA2 reversion

To establish that PARPi resistance is mediated by BRCA2 in our system and to directly assess how BRC-repeat retention influences revertant fitness, 12 reversion clones were isolated and genotyped from the p.S1982Rfs*22 background (Supplemental Fig. 7A). Next, a reversion clone derived from each of the parental alleles p.Y1569Rfs*2 (clone 1, splicing-mediated restoration resulting in loss of at least four BRC repeats) and p.S1982Rfs*22 (clone 6, splicing-mediated restoration resulting in loss of at least seven BRC repeats), was re-edited with a secondary sgRNA near the PALB2 interaction domain in exon 3 to test for the presence of a *BRCA2* variant with residual function in these clones. Following one week of passaging, cells were treated with either mock medium or 1 µM niraparib. NGS analysis showed that newly generated frameshift alleles were significantly depleted in both clones under niraparib treatment (Supplemental Fig. 7B). This contrasts with the parental p.S1982Rfs*22 frameshift clone, which was not sensitive to re-targeting in exon 3 (Fig. 1D). Interestingly, frameshift depletion was more pronounced in the p.Y1569Rfs*2 revertant which likely retains four BRC repeats than in the p.S1982Rfs*22 revertant which likely retains one, suggesting that the extent of BRC-repeat retention contributes to the relative fitness of PARPi-resistant reversions.

Nanopore sequencing of PARPi-resistant populations in both the p.Y1569Rfs*2 and p.S1982Rfs*22 genotypes revealed transcripts lacking all eight BRC repeats, prompting us to test whether these isoforms alone could mediate PARPi-resistance, or whether they arose as a byproduct of alternative splicing. To directly compare the fitness of a *BRCA2* frameshift allele to that of an in-frame *BRCA2* isoform lacking all BRC repeats, we generated an out-of-frame BRC deletion mutant (ΔBRC parental) in the p.S1982Rfs*22 background and edited the clone with sgRNA #392, which cleaved 9 bp downstream of the deletion (Fig. 7A). This design allowed direct comparison of the fitness of the parental ΔBRC frameshift allele and secondary in-frame ΔBRC restoration alleles generated by re-editing, as both could be quantified within the same NGS amplicon. Following re-editing, cells were passaged for one week before being treated with either mock or 0.25 µM, 0.5 µM, or 1 µM niraparib media. AP staining showed no increase in survival of the re-edited population relative to the non-edited ΔBRC parental control following niraparib treatment (Supplemental Fig. 8A, B). Surviving populations from the mock and 0.25 µM niraparib conditions were subsequently analyzed by NGS. Notably, in-frame ΔBRC restoration alleles were depleted relative to the parental and compound frameshift alleles even under mock conditions, suggesting a growth disadvantage independent of PARPi treatment (Fig. 7B). Although less pronounced, this depletion persisted following niraparib treatment, potentially reflecting reduced cell proliferation under PARPi treatment. Together, these results indicate that restoration of the *BRCA2* reading frame in the absence of all eight BRC repeats is insufficient to confer PARPi resistance and may impose an additional fitness disadvantage relative to frameshift alleles.

**Figure 7:**
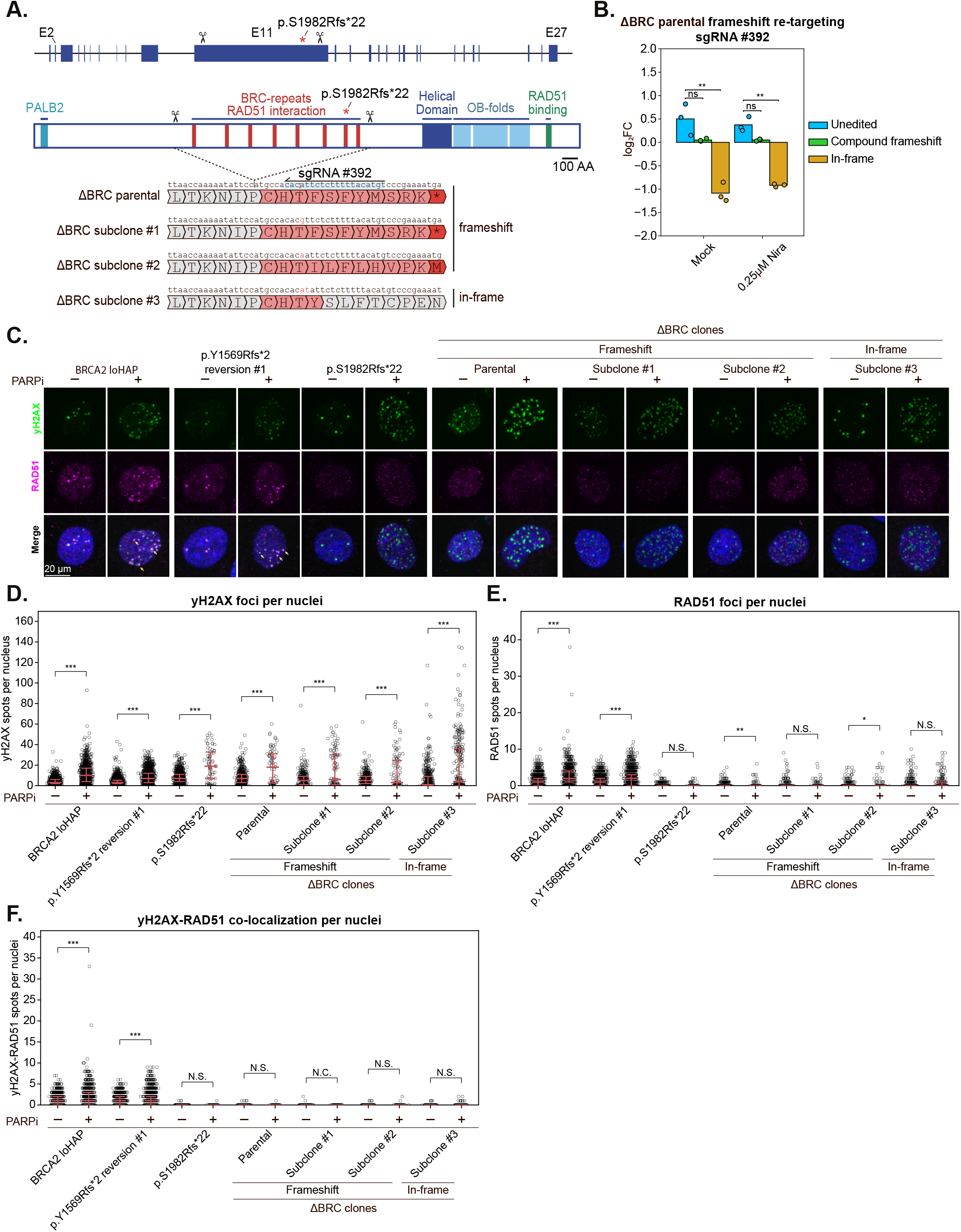
In-frame *BRCA2* restoration lacking all eight BRC repeats fails to confer PARPi resistance. (A) Schematic of *BRCA2* gene and protein architecture and the ΔBRC editing strategy. First, two sgRNAs were used to generate an out-of-frame deletion of the BRC repeats in the p.S1982Rfs*22 parental cell line. This ΔBRC parental cell line was then re-edited with sgRNA #392 to restore the reading frame of the protein. Three subclones were isolated following re-editing, including two compound frameshifts (ΔBRC subclones #1 & #2) and an in-frame subclone, ΔBRC #3. Representative translations are shown for each genotype along with the genetic changes. Exact deletion coordinates can be found in the supplementary data. (B) Log2 fold change of unedited, compound frameshift, and in-frame ΔBRC alleles following re-targeting of the ΔBRC parental frameshift line with sgRNA #392 and treatment with mock medium or 0.25 μM niraparib relative to day 7. Inframe ΔBRC alleles were not enriched under niraparib treatment. No survivors were observed at higher niraparib concentrations. Statistical significance was assessed using an unpaired two-sided Welch’s t-test comparing replicate-level log2 fold-change values between unedited and mutant allele classes within each condition. (C) Representative immunofluorescence images of γH2AX (green) and RAD51 (magenta) in *BRCA2* loHAP cells, p.Y1569Rfs*2 reversion clone 1, the parental p.S1982Rfs*22 frameshift line, the ΔBRC parental frameshift line, two compound frameshift ΔBRC subclones, and the in-frame ΔBRC subclone #3 following mock or 0.5 μM niraparib treatment for 24 hours. Scale bar, 20 μm. Representative γH2AX-RAD51 colocalized foci are indicated by yellow arrows in PARPi-treated *BRCA2* loHAPs and p.Y1569Rfs*2 reversion clone 1. (D) Quantification of γH2AX foci per nucleus for the indicated genotypes under mock and PARPi treatment. (E) Quantification of RAD51 foci per nucleus for the indicated genotypes with no threshold. (F) Quantification of γH2AX-RAD51 colocalized foci per nucleus. For panels D–F, individual points represent nuclei, and horizontal lines indicate the median and interquartile range. While all nuclei are plotted, only nuclei with one or more foci were used to calculate statistical significance. Statistical significance is indicated as follows: not significant (N.S.), not calculated (N.C.), not detected (N.D.), ***P < 0.001, **P < 0.01, *P < 0.05.

To further characterize the ΔBRC *BRCA2* isoform, two compound frameshift subclones and one in-frame ΔBRC subclone were isolated from the re-edited ΔBRC parental population (Fig. 7A). Each of these subclones, along with unedited *BRCA2* loHAPs, p.Y1569Rfs*2 reversion clone 1, p.S1982Rfs*22, and the parental ΔBRC frameshift cell line where independently assayed for niraparib sensitivity with AP staining and for and for HR activity by immunofluorescence analysis of RAD51 and γH2AX foci^13,41^. Consistent with our previous results, AP staining showed that both the in-frame ΔBRC subclone #3 and all out-of-frame cell lines were sensitive to niraparib treatment relative to the p.Y1569Rfs*2 reversion clone 1 and the unedited *BRCA2* loHAPs (Supplemental Fig. 8C - E). Immunofluorescence analysis of PARPi-treated cells revealed an increase in γH2AX-marked DNA damage foci across all genotypes relative to mock-treated controls (Fig. 7C, D). In contrast, only the BRCA2 loHAPs and p.Y1569Rfs*2 reversion clone 1 showed a robust increase in RAD51 foci upon PARPi treatment (Fig. 7C, E). RAD51 foci were detectable in the in-frame ΔBRC and frameshift subclones, yet they appeared diffusely distributed throughout the nucleus and did not colocalize with γH2AX-marked DNA damage foci (Fig. 7C, F). In contrast, both unedited BRCA2 loHAPs and p.Y1569Rfs*2 reversion clone 1 exhibited significant increases in RAD51–γH2AX colocalization following PARPi treatment (Fig. 7C, F). Together, these results suggest that restoration of the BRCA2 reading frame in the absence of all eight BRC repeats is insufficient to confer PARPi resistance and support efficient RAD51 recruitment to sites of DNA damage.

## Discussion

PARPi-resistant *BRCA2* reversions are clinically important, yet the molecular constraints that govern which restoration events arise and which ultimately confer resistance remain poorly defined. Here, using an isogenic cell system combined with targeted genome editing, pooled CRISPR mutagenesis, and long-read transcript analysis, we define the principles governing *BRCA2* reversion and demonstrate that reversion patterns follow defined principles and are therefore predictable.

Reversion mutations occur most frequently near the primary mutation and decline with increasing distance between the frameshift and secondary mutation. This relationship is further shaped by the local architecture of PTCs, which restricts the spectrum of secondary mutations that can restore the reading frame. These observations provide a mechanistic explanation for the tendency of clinical *BRCA2* reversions to cluster within 100 bp of the pathogenic mutation^17^, are recapitulated in clinical data^33^, and suggest that reversion risk is partially encoded in the local sequence context of each pathogenic allele. While local sequence context dictates the spectrum of available restoration mutations, it does not predict which restoration events will be selected as PARPi-resistant reversions. *BRCA2* restoration events are further selected at the regional level to preserve essential functional domains. This principle provides a mechanistic explanation for the clinically observed C-terminal reversion cold spot^17,31,32^. Unlike retrospective clinical analyses, our system enables detection of restoration events both before and after PARPi selection, providing a means to distinguish events that are generated from those that are subsequently selected. The recurrence of these patterns in an isogenic background indicates that they are intrinsic features of *BRCA2* reversion, rather than the result of variations in tumor type, treatment history, or genetic background. Thus, functional selection is superimposed on local sequence determinants, such that only a subset of restoration events that preserve essential BRCA2 domains confer PARPi resistance.

Interexonic restoration events followed the same general logic as local intraexonic events: their formation was constrained by the proximity and distribution of PTCs, whereas their ability to confer PARPi resistance was filtered by preservation of essential BRCA2 domains. Alleles with extended neopeptide tails spanning exon boundaries are therefore more permissive to distal interexonic restoration than alleles containing proximal PTCs. Consistent with this model, the p.S158Vfs*24 and p.P143Gfs*21 alleles preferentially generated PARPi-resistant reversions involving exons 6 and 7, whereas p.D156Efs*12 favored exon 6 only, reflecting the distinct downstream PTC landscapes created by each frameshift. The p.P143Gfs*21 mutation also altered local splicing and produced a reversion pattern resembling that associated with the clinical splice-site variant c.475G>A^31^. The failure of exon 2/3 and exon 23/24 restoration events to confer resistance further shows that restoring an open reading frame alone is insufficient when essential BRCA2 functions are disrupted. Conversely, the presence of revertants lacking exons 5, 6, 7, or 13 indicates that these regions are dispensable for PARPi resistance. Together, these findings extend previous observations of interexonic reversions in clinical samples and directly define the sequence and functional constraints governing their formation.

The exon 11 reversion spectrum was unexpected on several levels. Exon 11 was permissive to unusually large restoration events. However, these events were not unrestricted and remained primarily confined to exons 11 and 12, recapitulating clinically observed patterns of exon 11 reversion. In some cases, these events resulted in splicing-mediated isoforms that remove all eight BRC repeats. This mechanism was not random. Long-read sequencing further showed that these restoration events arose through recurrent cryptic splice-donor usage. Nearly all high-probability in-frame cryptic donor sites were used to generate splicing-mediated restoration events, whereas the use of predicted in-frame cryptic acceptors was not detected. Importantly, three of the five isoforms detected here were shared across multiple pathogenic frameshift alleles, including the clinical variant p.S1982Rfs*22. These findings indicate that splicing-mediated restorations in exon 11 are directionally biased, restricted to a narrow set of donor sites, and follow a reproducible pattern. Furthermore, they suggest that a broad range of pathogenic mutations could be restored through the use of several cryptic splice donors, highlighting the need for transcript-level surveillance of a small set of recurrent isoforms in clinical settings.

A major advantage of the *BRCA2* loHAP system is that reversions can be evaluated in a genetically defined, isogenic background in which copy-number profiling confirms the presence of a single *BRCA2* allele^21^. This allows the contribution and relative fitness of individual reversion genotypes to be assessed without ambiguity from a second *BRCA2* copy. Our data suggest that this fitness is graded rather than binary: for example, the exon 11 revertant retaining four BRC repeats showed a stronger fitness advantage than the revertant retaining only one BRC repeat, consistent with a mouse study indicating reduced fitness of BRCA2 proteins containing a single BRC repeat^42^. However, conflicting data exist on whether direct BRCA2 reading-frame restoration events that delete all eight BRC repeats confer PARPi resistance and restore robust RAD51 recruitment^31,32^. In our cell system, a systematic head-to-head comparison of an in-frame allele lacking all BRC repeats with frameshift mutations in exon 11 did not reveal a growth advantage, PARPi resistance, or restoration of RAD51 foci. These findings support the conclusion that retaining at least one BRC repeat is required for functional reversion in our cell system.

Our results demonstrate that *BRCA2* reversion is governed by predictable sequence-based and regional constraints. Local sequence context determines which secondary restoration events can form, whereas protein domain architecture determines whether those mutations constitute PARPi-resistant reversions. This logic applies to restoration events occurring both within and between exons. Applied to exon 11, these principles suggest that a single BRC repeat is sufficient for PARPi resistance. These findings have direct clinical implications for identifying allele-specific regions at elevated risk of reversion, interpreting the functional consequences of secondary mutations, and determining which pathogenic *BRCA2* alleles are most likely to confer durable responses to PARPi.

Interpretation of the restoration patterns presented in this study should account for the fact that these events were generated using the CRISPR-Cas9 system. Although local sequence context and microhomology strongly influence repair of DSBs induced by both Cas9 and endogenous mechanisms^29,30^, the resulting mutation spectra may differ. Cas9-mediated repair in our system predominantly generates insertions and deletions. Consequently, this approach may not capture the full spectrum of clinically occurring *BRCA2* reversions, including precise restoration to the wild-type sequence or nucleotide substitutions that restore BRCA2 function^17,43,44^. Precise restoration may be further underrepresented in the loHAP background because these cells contain only a single *BRCA2* allele and therefore lack an intact second copy that could serve as a homologous repair template.

## Material and methods

### Pluripotent stem cell culture

Human pluripotent stem cell research was approved by the Stem Cell Research Oversight Committee at the University of California, Berkeley (protocol 2012-12-024). WIBR3 hESCs^22^ (NIH stem cell registry no. 0079) were cultured on irradiated mouse embryonic fibroblasts (MEFs; 4.1 × 10^5^ cells cm^−2^) in hESC media composed of DMEM/F12, 20% KnockOut serum replacement, 1× non-essential amino acids (NEAA), 1 mM glutamine, 1× penicillin–streptomycin, 0.1 mM β-mercaptoethanol, and 4 ng ml^−1^ heat-stable basic fibroblast growth factor. Media was changed daily. Cells were passaged every 5–7 days with 1 mg ml^−1^ collagenase IV, and 10 µM Y27632 (CD0141, Chemdea) was added on the day before and after single-cell passage to improve survival.

### Generation of *BRCA2* loHAP frameshift cell lines

Unedited *BRCA2* loHAP cells^21^ were nucleofected with Cas9–sgRNA ribonucleoprotein complexes containing a single sgRNA, as described below. After nucleofection, cells were plated directly onto three 96-well MEF feeder plates at densities of 10, 20, and 30 cells per well to account for batch-to-batch variation in cell survival. Media was changed on days 4, 7, 10, 12, and 13, and 10 µM Y27632 was added on day 13. On day 14, cells were incubated with 40 µL of 0.25% trypsin–EDTA for 5–10 min at 37 °C. Trypsin was neutralized by adding 60 µL of FBS/hPSC media supplemented with 10 µM Y27632, in which 10% KnockOut serum replacement was substituted for 10% FBS. Cells were gently triturated, and 50 µL of the resulting suspension was transferred to a 96-well PCR plate containing 50 µL of 2× lysis buffer (100 mM KCl, 4 mM MgCl₂, 0.9% NP-40, 0.9% Tween-20, and 500 µg ml^−1^ proteinase K in 20 mM Tris, pH 8.0). The remaining 50 µL was reseeded into a fresh MEF-coated 96-well plate preloaded with 100 µL of FBS/hPSC media supplemented with 10 µM Y27632 and cultured for an additional 7 days, with daily media changes. Lysates were incubated at 50 °C overnight and then at 95 °C for 10 min to inactivate proteinase K. Each well was genotyped by targeted NGS across the intended editing site.

### PARP inhibitor treatment and selection

The relative fitness of clonal *BRCA2* frameshift cell lines under PARP inhibitor selection was assessed by mixing a frameshift cell line with unedited *BRCA2* loHAPs at a 1:1 ratio and culturing the mixed populations for 1 week in mock media or media containing 0.5 μM or 1 μM niraparib before genomic DNA extraction. All experiments were performed with three biological replicates unless otherwise specified. A non-Cas9 nucleofected parental frameshift cell line was treated with PARPi in parallel with each experiment to account for spontaneous reversions; only experiments in which all control cells were killed following PARPi treatment were interpreted. Allele fitness under PARPi selection was inferred from changes in allele frequency measured by NGS of the region containing the primary frameshift allele. Relative depletion or enrichment of mutant and unedited alleles was calculated as the log2 fold change in allele frequency in the drug-treated population relative to the mock-treated population. To enrich for reversion events, cells were treated with 1 μM niraparib, unless otherwise indicated, or mock media for 5–10 days, with drug added 48 h after passaging.

### Nucleofection

hESCs cultured on MEFs were released from feeder cells by incubation with 1 mg ml−1 collagenase IV and 0.5 U ml^−1^ dispase for 20–25 min. Detached colonies were washed twice with DMEM/F12 and then dissociated into single cells by treatment with 0.25% trypsin (25200114, Thermo Fisher) for 5–10 min at 37 °C, followed by neutralization with hESC media. Cells were counted, washed once with 1× PBS, and 0.5–1 × 10^6^ cells were used per nucleofection, depending on expected survival. Cell pellets were resuspended in 20 µL Lonza P3 Primary Cell Nucleofector solution and mixed with either 80 pmol Cas9 protein (QB3 MacroLab, UC Berkeley) alone or preassembled Cas9–sgRNA ribonucleoprotein complexes (200 pmol sgRNA), with or without 100 pmol ssODN HDR donor. Nucleofection was performed using the program CA137 on a Lonza 4D-Nucleofector. The nucleofected cells were divided into three biological replicates that were maintained independently for all experiments.

### Dual-editing strategy for HDR frameshift alleles

Nucleofections were conducted as described above. Designer frameshift alleles were introduced via 100–150 nt ssODN HDR templates centered on the cleavage site of the first sgRNA. HDR templates were designed with synonymous CRISPR–Cas9-blocking mutations in either the PAM or the 3’ end of the first sgRNA; synonymous alleles were also introduced in all PTCs in frames 2 and 3, located between the cut site of the first and second sgRNA. The second sgRNA was not blocked from cutting the HDR template. HDR templates contained a ≥2 nucleotide change relative to the wild-type sequence to distinguish designer mutations from sequencing errors. 100 pmol of the ssODN library (purchased from Integrated DNA Technologies (IDT)) was co-delivered with the first sgRNA in a preassembled Cas9 RNP into 1 million cells. Edited cells were divided into three biological replicates, cultured for one week, singularized, and pooled before re-editing another 1 million cells with the second sgRNA in a Cas9-RNP mixture. The cells were then seeded into three wells as biological triplicates and passaged with 0.25% trypsin–EDTA every 7 days for two weeks, before treatment with either mock or 1 µM niraparib. A cell pellet was collected at each time point and condition, and the pellet was sequenced in triplicate using an NGS amplicon encompassing the region with the restoration mutation. Alleles were called and quantified by CRISPResso2. Restoration alleles were distinguished from single, in-frame edits using neopeptides unique to each locus. In-frame alleles are those that contain large deletions encompassing both sgRNA cut sites that cannot be confidently identified as a restoration event, dual in-frame edits by both sgRNAs, and single in-frame edits by either the first or second sgRNA. HDR templates can be found in the supplementary materials.

### PCR preparation

DNA was extracted from each time point or drug treatment condition using phenol-chloroform DNA extraction. Each PCR was conducted in triplicate, amplified using GoTaq polymerase (M300A, Promega), and contained 300 ng of DNA in a 10 µL volume. PCR quality was confirmed with gel electrophoresis prior to NGS. Amplicons were purified and processed according to previously described methods^45^.

### NGS analysis and quantification of reversion frequencies

FASTQ files were initially analyzed with CRISPResso2^46^ to merge paired-end reads, perform quality filtering, and generate allele-frequency tables. These allele-frequency tables were then processed with a custom Python script for reversion allele classification. Briefly, aligned amplicon sequences were translated, merged across technical PCR replicates, and collapsed into unique alleles. Translation was performed using locus-specific 5′ and 3′ sequence boundaries, which were defined according to the position of the primary frameshift mutation and the secondary sgRNA target site. Alleles were then normalized to total read depth to generate allele frequencies and compared with the corresponding week 1 samples to calculate fold change. Heat maps, dot plots, and other visualizations were generated using Matplotlib and Seaborn. Allele frequencies used to generate visualizations are listed in the supplementary data files. Custom scripts for quantifying and visualizing allele frequencies are available on the Hockemeyer lab GitHub.

For experiments performed in clonal frameshift cell lines, restoration alleles were defined as any translated allele within the analyzed region that restored the *BRCA2* reading frame. Compound frameshift alleles were defined as parental frameshift alleles that acquired a secondary edit but did not restore the reading frame. “Other” alleles contained mutations or deletions that altered or removed the 5′ or 3′ sequence boundaries required for translation calling in the custom script. Restoration alleles were classified as genetic if DNA-level insertions or deletions restored the *BRCA2* reading frame while preserving canonical splice junctions. Alleles were classified as splicing-mediated if restoration of an in-frame *BRCA2* transcript required altered splice-site usage, including exon skipping or engagement of cryptic splice donors or acceptors. Alleles were categorized as intraexonic or interexonic on the basis of whether the restoring event occurred within the exon containing the primary frameshift mutation or extended across exon boundaries. Where applicable, restoration alleles were further distinguished from single in-frame editing events by the presence of locus-specific neopeptides unique to each engineered restoration interval. All allele frequencies used to generate the figures are available in the supplementary materials.

sgRNA frequencies were assessed by NGS of the lentiviral backbone and processed as above, except for allele translation. The cDNA cut position for each sgRNA was derived from the start of translation relative to the *BRCA2* transcript isoform 1 (MAVE transcript ID: ENST00000380152.8).

### Lentiviral sgRNA design, cloning, and infection

sgRNA libraries were cloned into the pGuide backbone (Addgene plasmid #64711) using an ssODN library purchased from IDT. The pGuide vector was digested with FastDigest Esp3I (FD0454, Thermo Fisher), and the ssODN library was assembled into the backbone using NEBuilder HiFi DNA Assembly (E5520S, NEB). Lentivirus was produced from the pooled plasmid library and used to infect one confluent six-well plate of cells seeded on puromycin-resistant MEF feeder plates at a multiplicity of infection of approximately 0.1. Infected cells were cultured for 48 h before selection in hESC media containing 0.8 μg ml^−1^ puromycin. Selection was continued for at least one week, or until nucleofection. Transduced cells were then nucleofected with 80 pmol Cas9 protein in 10 µL water per reaction. Each reaction contained 2 × 10^6^ cells. Between one and three nucleofections were conducted per experiment, depending on the scale of the sgRNA library. Sequences of sgRNAs used to generate the libraries are listed in the supplementary data table.

### RNA isolation, cDNA synthesis, and transcript analysis

RNA was isolated using the RNeasy Mini Kit (74104, Qiagen) with on-column DNase treatment where indicated. cDNA was synthesized using the ProtoScript II Reverse Transcriptase kit using the standard protocol with d(T)23VN primers (M0368, NEB). Regions spanning candidate interexonic or splicing-mediated restoration events were amplified using the specified primers in flanking exons, with Q5 polymerase in 25 µL reactions containing 2 µL of cDNA diluted to 1:50 (M0493S, NEB). PCR products were analyzed by agarose gel electrophoresis and purified with the Monarch Spin High-Capacity DNA Cleanup Kit (T1135L, NEB). PCR products were analyzed with either NGS, Sanger sequencing, or long-read nanopore sequencing.

### Long-read nanopore sequencing

To resolve complex exon 11 restoration events, cDNA was synthesized as described above from 1 μM niraparib-treated edited populations of the p.Y1569Rfs*2 and p.S1982Rfs*22 clones, together with the corresponding unedited parental lines. Two PCR amplicons were generated to capture exon 11 restoration products: (1) a forward primer spanning the exon 10/11 boundary paired with a reverse primer in exon 12, and (2) a forward primer in exon 10 paired with a reverse primer spanning the exon 11/12 boundary. PCR products were amplified and purified as described above. 40 ng of PCR products were barcoded with the Nanopore Native Barcoding library kit (SQK-NBD114.24) per the manufacturer’s steps, with the following modifications. DNA fragment end-prep was performed without the DNA Repair component in 1.75 µL of NEBNext Ultra II End Prep Reaction Buffer, 0.75 µL of End Prep Enzyme Mix in a 15 µL reaction volume. DNA fragments were cleaned up with a 1.5x ratio of Pronex beads (Promega) instead of AMPureXP beads. Bead drying after pooling of barcoded DNA fragments was extended to 30 minutes as required to ensure no residual ethanol carryover prior to Native Adapter (NA) ligation. For the last step, Native Barcoding DNA fragments were eluted from AMPureXP beads in EB buffer after SFB buffer washes according to the manufacturer’s protocol. The entire library (5-9 barcodes) was sequenced on a MinION flow cell (FLO-MIN114), yielding at least 300k reads per barcode. Nanopore reads were aligned to the wild-type *BRCA2* cDNA (MAVE transcript ID: ENST00000380152.8) reference using minimap2 v3^47^. Alignments were visualized in IGV^48^, and deletion breakpoints were resolved for each allele. Genetic restorations were defined as events that restored the reading frame within exon 11 while retaining canonical splice junctions. Splicing-mediated restoration events were defined as events with a 5′ breakpoint in exon 11 and a 3′ breakpoint at exon 12, or with a 5′ breakpoint at exon 10 and a 3′ breakpoint within exon 11.

### Alkaline phosphatase staining

Cells were cultured for between 10 and 14 days under mock or drug treatment before alkaline phosphatase (AP) staining according to a previously described protocol^49^. Images were quantified in ImageJ by measuring mean staining intensity for each well. Relative survival was calculated by normalizing the mean intensity of drug-treated wells to the corresponding mock-treated control.

### Immunofluorescence Staining

Cells were plated on Matrigel (Corning) coated Phenoplate-96 imaging plates (Revvity) at a seeding density of 10,000 cells per well 24 hours prior to fixation with 4% Paraformaldehyde for 10 minutes at room temperature. Following a wash in 1x PBS, cells were permeabilized in 0.1% Triton X-100 for 15 minutes at room temperature, before an additional wash in 1x PBS and 15 min blocking in Blocking buffer (3% BSA, 5% horse serum, 0.1% Tween-20 in PBS) at room temperature. Primary (anti-RAD51: 1/2,000, anti-γH2AX: 1/2,000) and secondary antibody (Alexa 488-Anti-rabbit IgG: 1/5,000, Alexa 546-Anti-mouse IgG: 1/5,000) staining occurred consecutively at room temperature for 1 hour each. All antibodies were diluted in blocking buffer as described above. Nuclei were stained with 0.05 µg/mL DAPI (Sigma-Aldrich) in PBS, for 15 minutes at room temperature. Cells were washed twice in 1x PBS with 0.1% Tween-20 after each step of staining: primary, secondary, and DAPI.

All immunofluorescent images were captured using the Opera Phenix high-content imager with a 63x objective and analysis was done using Harmony software (Revvity) on maximum-projected z-stacks. TIFs were called based on >50% overlap of foci.

## Acknowledgements

We thank Katherine Nathanson for advice on the experimental design and interpretation of BRCA2 reversion mutations. Johannes Schabort and Shayan Hosseinzadeh for critical comments on the manuscript. The Hockemeyer lab is supported by the Innovative Genomics Institute, Siebel Stem Cell Institute, the American Cancer Society (133396-RSG-19-029-01-DMC), and the NIH R01HL131744-05 and R21AG095841-01. A.H. is funded by the Li lab, which is supported by the Innovative Genomics Institute, CRISPR Cures for Cancer Initiative, and Shanahan Family Foundation. The California Cancer Research Coordinating Committee (CRCC) has funded tuition for the 2025-6 academic year for A.H.

## Author contributions

A.H., H.L., and D.H. conceptualized the project and designed the experiments. A.H. performed all experiments with the support of S.T., I.P.L., C. S., and F.L.K.. R.L. performed the nanopore sequencing of *BRCA2* cDNA. A.H. and D.H. analyzed the data and wrote the manuscript.

