## Supplemental_figures for "Decoding the *BRCA2* reversion principles underlying PARP inhibitor resistance"

### Horacek et al. Supplemental Figure 1

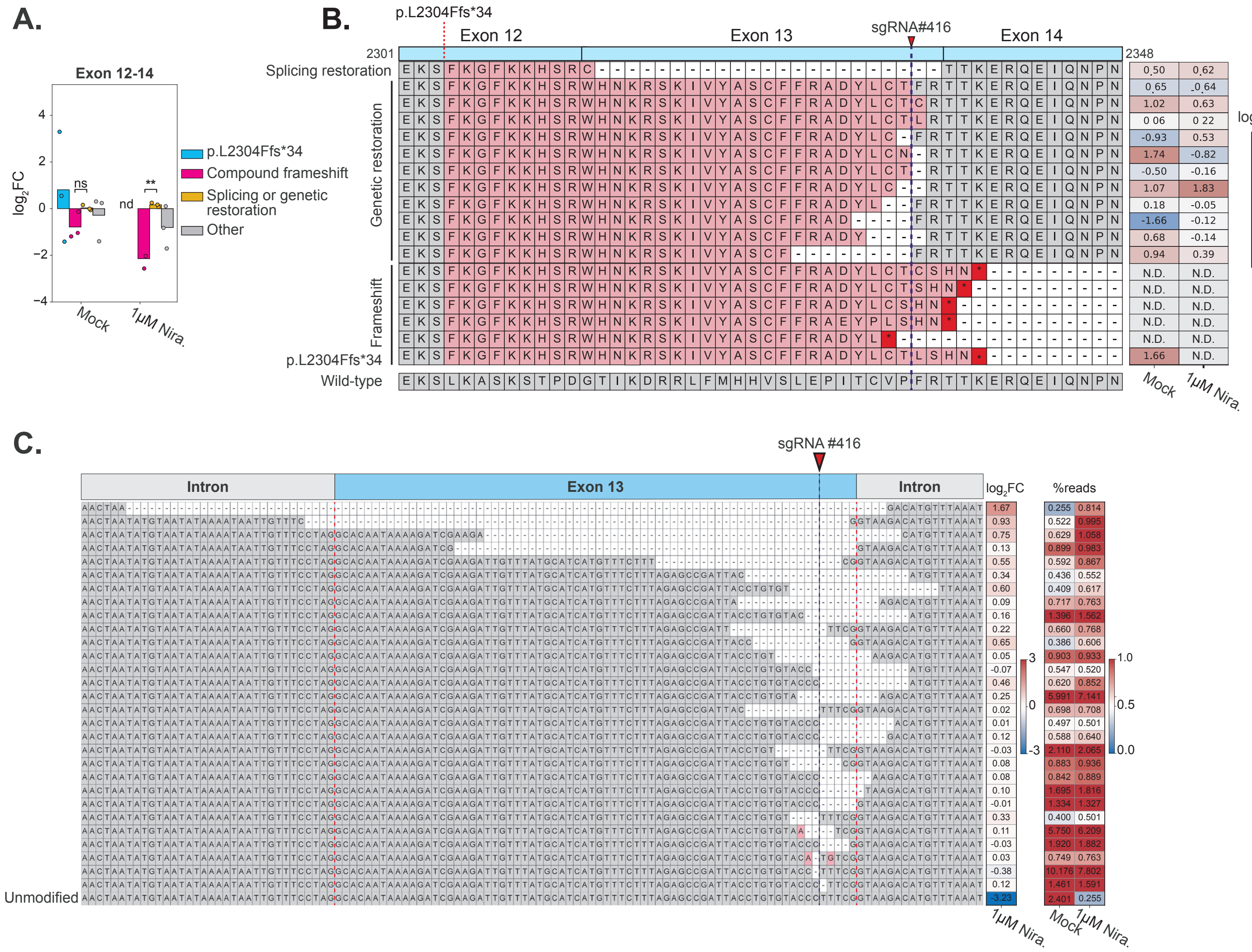

**Supplemental Figure 1: Interexonic restorations arise through distinct mechanisms.** (A) Average log2 fold change of alleles in *BRCA2* exons 12-14 at the cDNA level in mock and 1 µM niraparib conditions relative to day 7. Allele frequencies were obtained after nucleofecting the p.L2304Ffs*34 parental cell line with a single sgRNA targeting the 3' end of exon 13. (B) Representative translated alleles recovered from the p.L2304Ffs*34 background following editing at the exon 13/14 junction. Two classes of interexonic restoration alleles were identified: DNA-level (genetic) restorations that restored the reading frame across exons 12–14, and likely splicing-mediated restorations generated by exon 13 skipping, although large deletions removing exon 13 cannot be ruled out. Neopeptide sequences are highlighted in red. The sgRNA cut site is indicated above. Values in the right panel show average log2 fold change in the mock and 1 µM niraparib conditions relative to day 7. A Welch's t-test comparing the average replicate log2 fold change between p.L2304Ffs*34 and mutant allele types within each condition was used. Statistical significance is indicated as follows: non-significant (NS), not detected (ND), ***$P < 0.001$, **$P < 0.01$, *$P < 0.05$. (C) NGS analysis of gDNA from the p.L2304Ffs*34 frameshift clone edited with an sgRNA targeting the 3' end of exon 13. The configuration of exon 13, the 5' and 3' introns, and the sgRNA cut position is shown above. The DNA sequence alignments are shown in the left panel, with the corresponding log2 fold change values relative to day 7 in the middle panel and the mean allele frequencies in the far-right panel. The intron/exon boundaries are indicated by red dashed lines. Deletions removing exon 13, either the canonical splice donor or acceptor, and those internal to the exon are observed.

### Horacek et al. Supplemental Figure 2

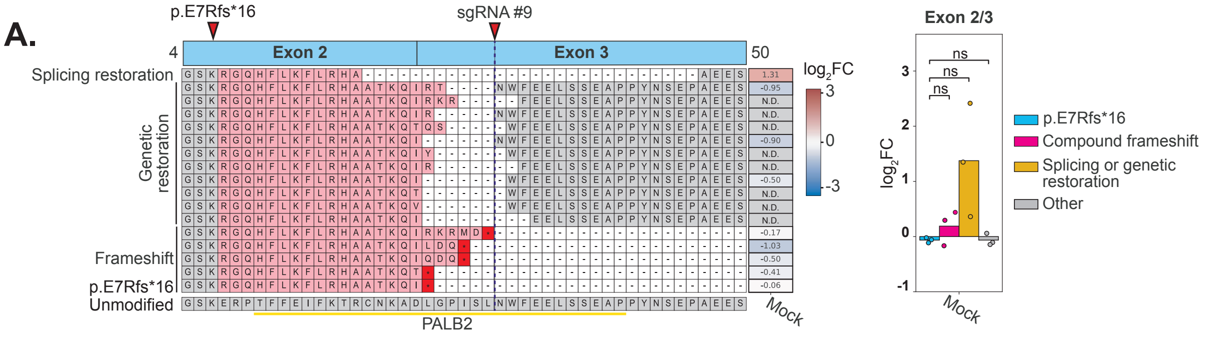


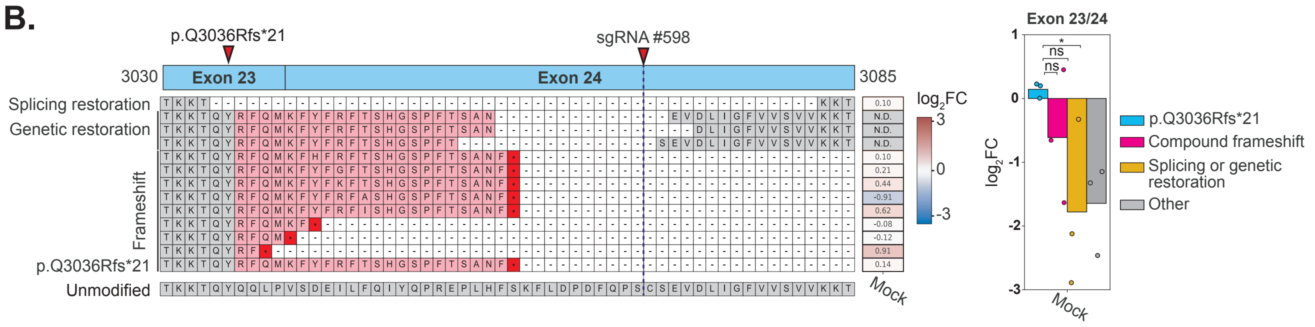

**Supplemental Figure 2: Interexonic restorations across key functional domains are not tolerated.** (A) NGS analysis of cDNA encompassing the exon 2-3 boundary. The exon 2 frameshift mutant (p.E7Rfs*16) was edited with a second sgRNA targeting exon 3. Restoration alleles that restored the reading frame were observed in the mock condition, but none survived treatment with 1 μM niraparib. Allele translations are shown on the left, with genetic and splicing restorations indicated. The PALB2 interaction domain is represented as a yellow line. (B) NGS analysis of cDNA obtained from the exon 23-24 boundary for the p.Q3036Rfs*21 clone targeted with an sgRNA in exon 24. Both exons are located within the BRCA2 OB folds. Splicing and genetic restorations are indicated. Log2 fold change values for both experiments are relative to day 7. All experiments were performed in triplicate. Statistical significance was assessed using an unpaired two-sided Welch's t-test comparing the average replicate log2 fold change between the parental frameshift and restoration/frameshift alleles within each condition. Statistical significance is indicated as follows: non-significant (NS), not detected (ND), ***$P < 0.001$, **$P < 0.01$, *$P < 0.05$.

### Horacek et al. Supplemental Figure 3

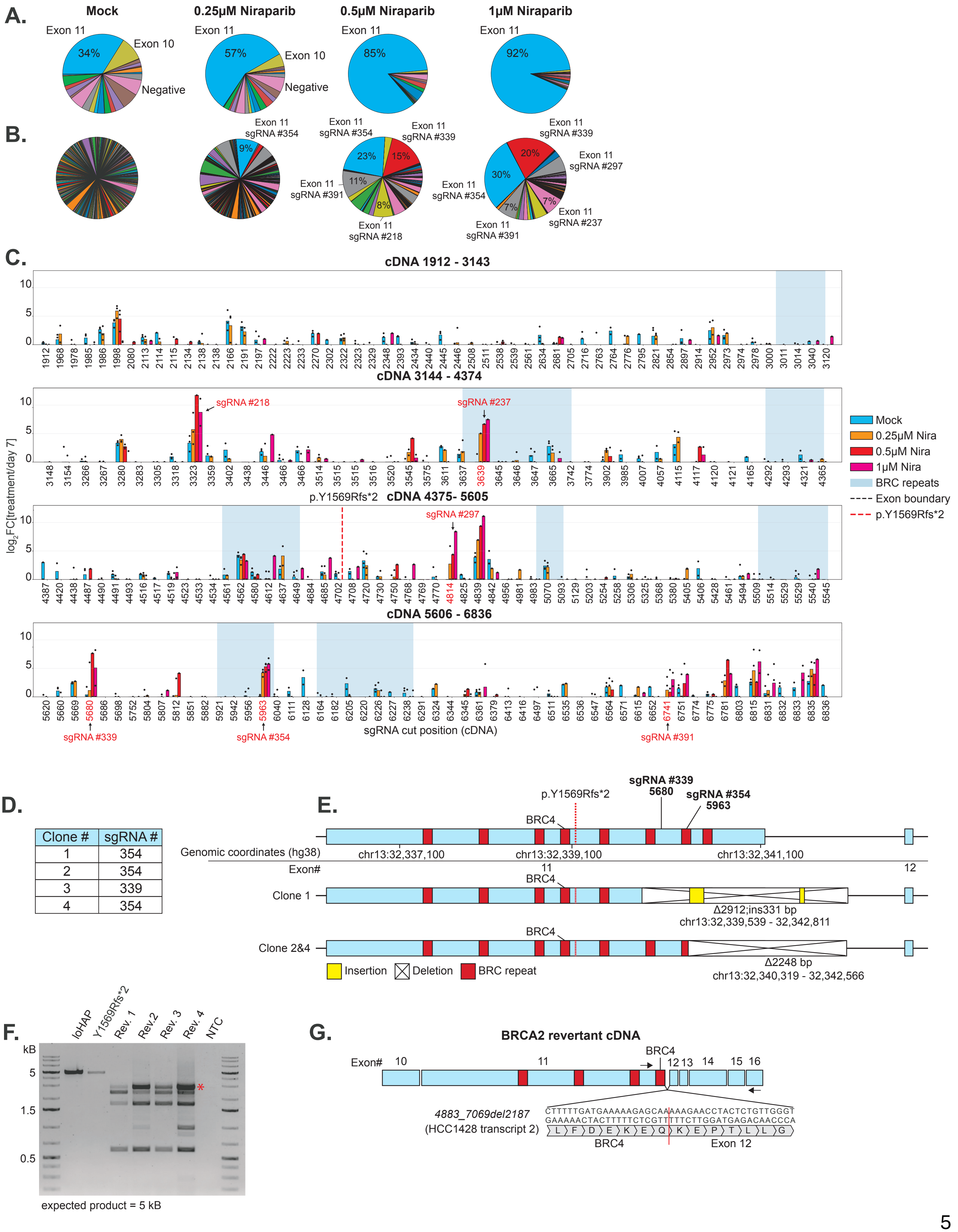


| Clone # | sgRNA # |
|---|---|
| 1 | 354 |
| 2 | 354 |
| 3 | 339 |
| 4 | 354 |

**Supplemental Figure 3: Exon 11 is permissive to large-scale *BRCA2* restoration events arising from cryptic splice donors.** (A) Pie charts showing average sgRNA frequency per exon across increasing niraparib concentrations (mock, 0.25 μM, 0.5 μM, and 1 μM) in the p.Y1569Rfs*2 background. As drug concentration increased, sgRNAs targeting exon 11 dominated the population, indicating enrichment of PARPi-resistant restoration events in this region. (B) Average distribution of enriched sgRNAs within exon 11 under PARPi selection. A small subset of sgRNAs (237, 297, 339, 354, and 391) accounted for a large proportion of the population at higher concentrations. (C) Average log2 fold change of sgRNA frequency across exon 11 based on sgRNA cut position for mock, 0.25 μM, 0.5 μM, and 1 μM niraparib concentrations relative to day 7 (n = 3 biological replicates). Enriched sgRNAs are labeled in red. BRC repeat regions and p.Y1569Rfs*2 are indicated. (D) sgRNA genotype present within each clone. (E) Genomic characterization of reversion alleles isolated from individual PARPi-resistant clones. Distinct alleles (Δ2912;ins331 bp and Δ2248 bp) were identified downstream of the primary mutation, spanning regions that include multiple BRC repeats. sgRNA positions and genomic coordinates (hg38) are indicated. (F) Agarose gel electrophoresis of cDNA PCR products amplified using primers that span from the exon 10-11 junction to exon 12 in revertant clones. The expected product size is 5 kB. A red asterisk indicates the largest conserved restoration isoform that was shared across all four clones. (G) Schematic of the structure of the largest conserved BRCA2 restoration isoform. The deletion junction of this isoform was amplified and sequenced using a forward primer near BRC repeat 4 and a reverse primer in exon 16. This restoration transcript arose through alternative splicing of a cryptic splice donor at the end of BRC4 and the canonical exon 12 splice acceptor, restoring the reading frame and deleting BRC repeats 5 - 8.

### Horacek et al. Supplemental Figure 4

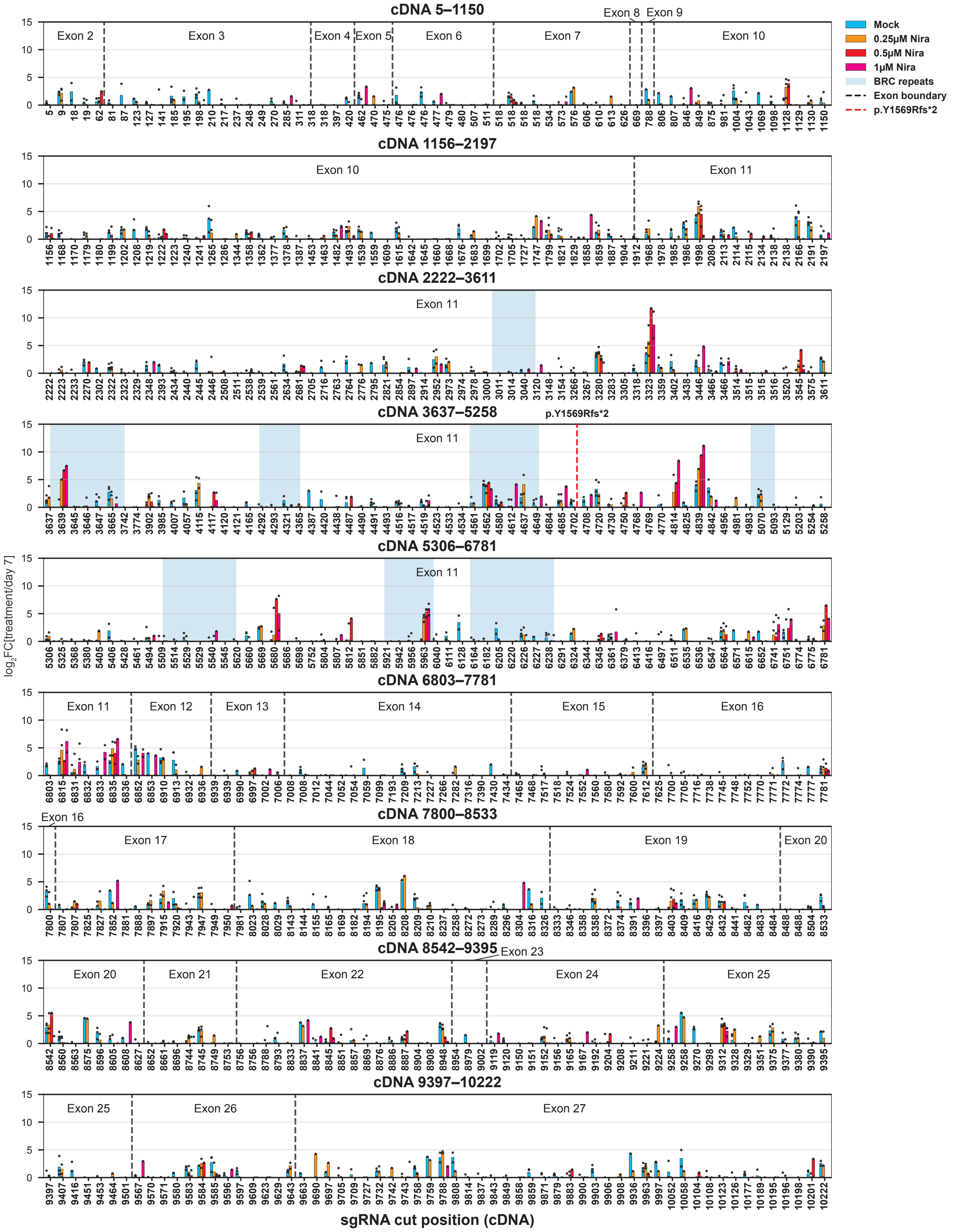

**Supplemental Figure 4: PARPi-resistant reversions in the p.Y1569Rfs*2 clone are confined to exons 11 and 12.** sgRNA enrichment, sorted by cut position, for the *BRCA2* cDNA-wide lentivirus experiment in the p.Y1569Rfs*2 (exon 11) frameshift clone. Exons are labeled, and their boundaries are shown as black dashed lines. The parental frameshift mutation is indicated by a red dashed line. BRC repeats are shaded blue. Average log2 fold-change sgRNA enrichment in mock (blue), 0.25 μM (orange), 0.5 μM (red), and 1 μM niraparib (pink) relative to day 7 (n = 3 biological replicates).

### Horacek et al. Supplemental Figure 5

**A.**

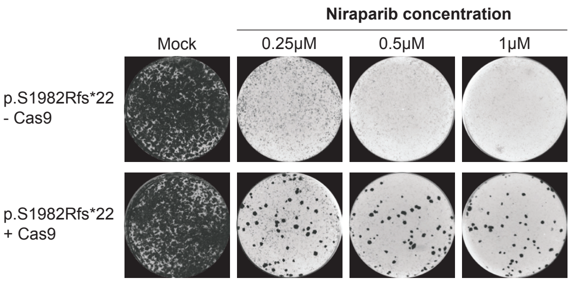


**B.**

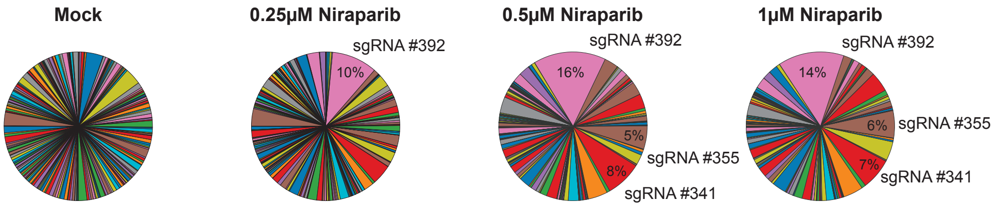


**C.**

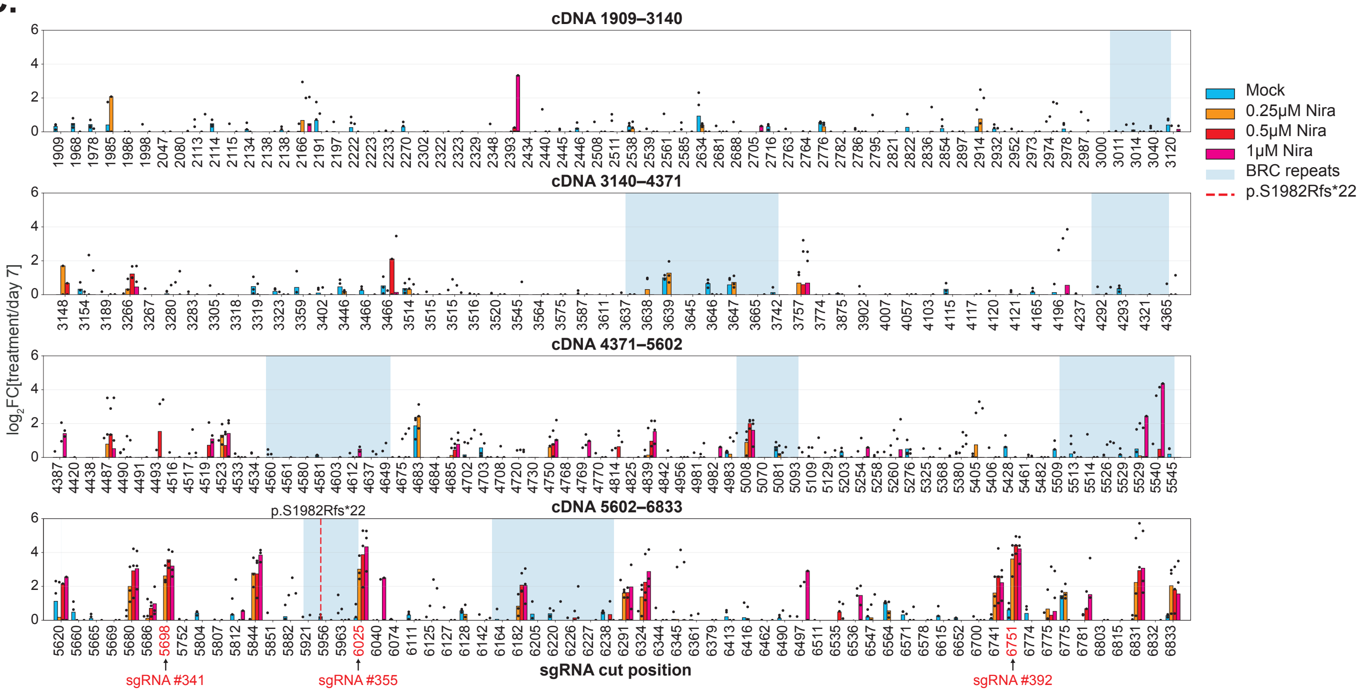

**Supplemental Figure 5: Personalized reversion map for the exon 11 *c.5946delT* (p.S1982Rfs*22) founder mutation.** (A) Alkaline phosphatase staining of pooled biological replicates of p.S1982Rfs*22 cells with or without Cas9, treated with increasing concentrations of niraparib (mock, 0.25 µM, 0.5 µM, and 1 µM). Colonies were observed only in Cas9-treated cells under niraparib selection, indicating the emergence of PARPi-resistant revertants. (B) Pie charts showing sgRNA representation across treatment conditions. A subset of sgRNAs, including sgRNA #341, #355, and #392, became enriched with increasing niraparib concentration. (C) Average log2 fold change in sgRNA abundance relative to day 7, plotted by cDNA cut position and stratified by niraparib concentration. Enriched sgRNAs are indicated in red. sgRNAs were generally enriched within 300 bp upstream and downstream of the c.5946delT mutation and within 100 bp of the 3' end of exon 11, with prominent peaks corresponding to sgRNAs #341, #355, and #392 (n = 3 biological replicates).

### Horacek et al. Supplemental Figure 6

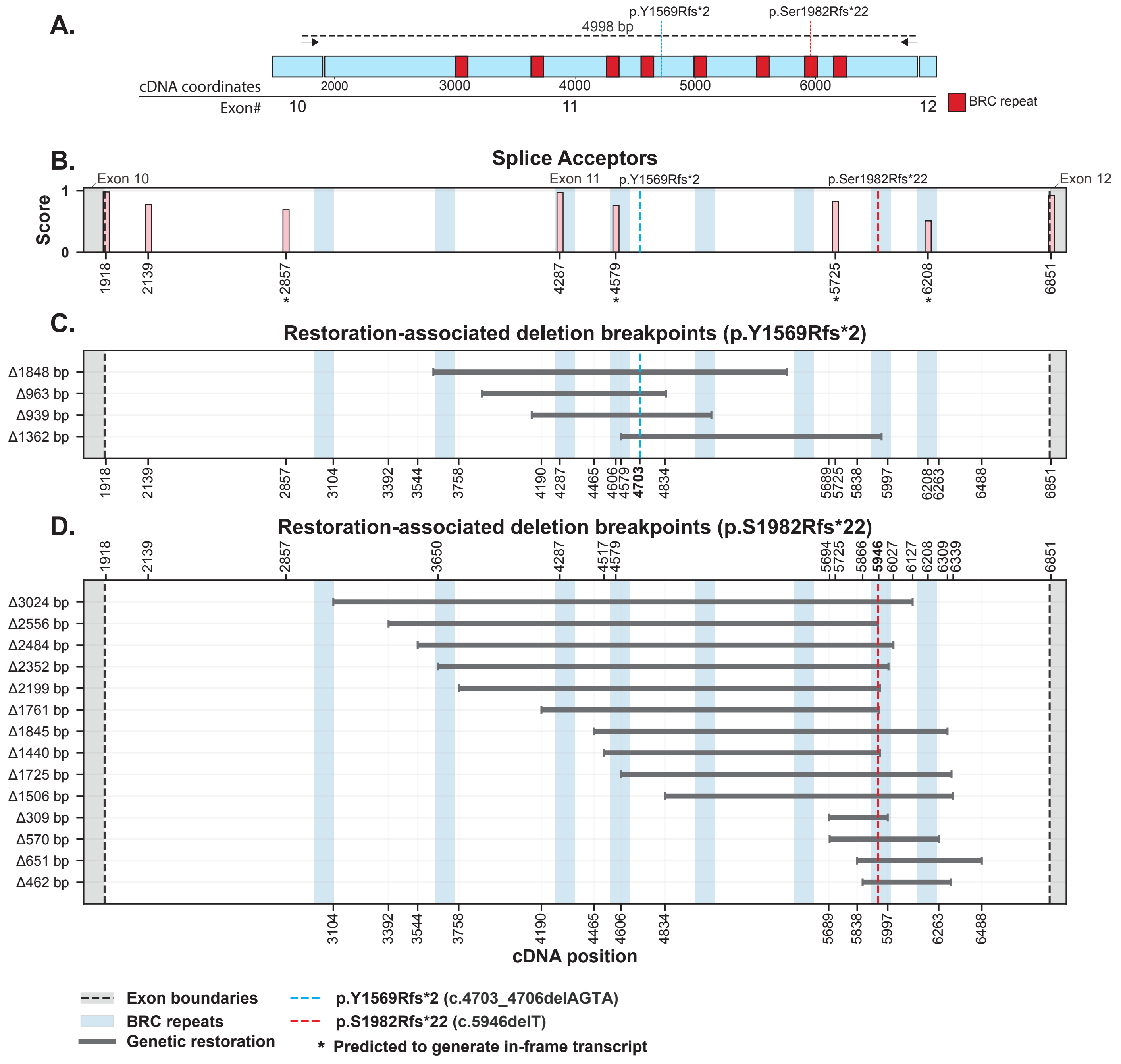

**Supplemental Figure 6: Alternative splice acceptors are not engaged in exon 11 splicing-mediated restorations.** (A) Schematic of the PCR strategy used for nanopore sequencing of *BRCA2* cDNA. PCR primers were anchored at exon 10 and at the exon 11-12 junction. The expected PCR product is 4998 bp. (B) Predicted splice acceptor strengths across exons 10-12, plotted as a function of cDNA position. Splice-site strength predictions were generated using NNSPLICE 0.9. Alternative splice acceptors that generate an in-frame product are indicated by an asterisk. Only two of the in-frame splice acceptors could be utilized to generate an in-frame product in these experiments. Deletion breakpoints identified by long-read nanopore sequencing of the cDNA amplicon isolated from the 1 μM niraparib-treated condition in the p.Y1569Rfs*2 (C) and p.S1982Rfs*22 (D) backgrounds. Genetic restorations are shown in gray, genotype-specific splicing-mediated restorations in green, and shared splicing-mediated restorations detected in both genotypes in red. Exon boundaries are indicated by gray shading and dashed lines, BRC repeats are shaded in blue, and the positions of the parental frameshift alleles are indicated. No splicing-mediated restorations were observed in either background.

#### Horacek et al. Supplemental Figure 7

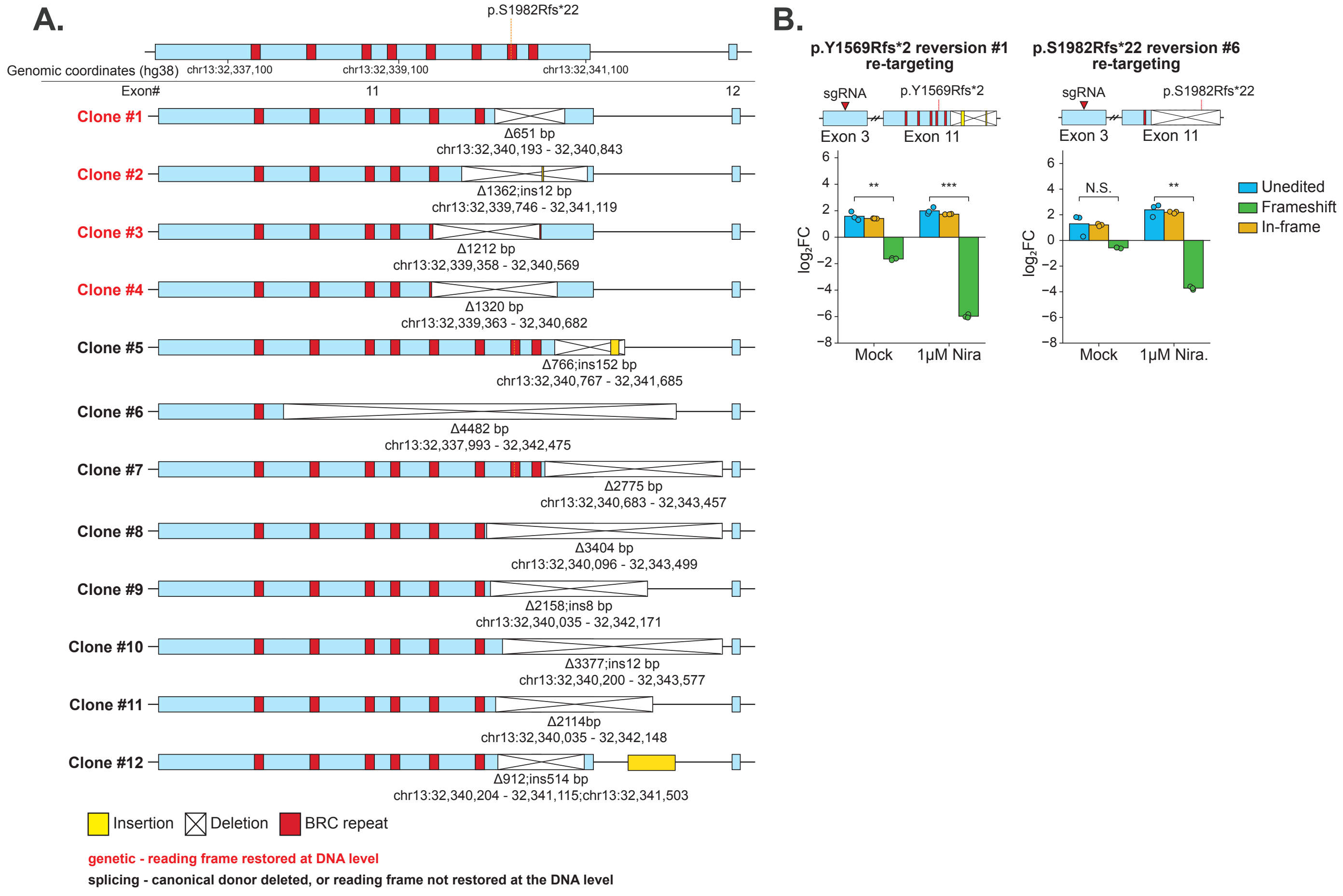

**Supplemental Figure 7: BRCA2 exon 11 reversions confer a graded fitness advantage.** (A) Genomic structures of 12 PARPi-resistant reversion clones isolated from the p.S1982Rfs*22 background. The position of the p.S1982Rfs*22 parental frameshift mutation is shown relative to *BRCA2* exon 11, exon 12, and the eight BRC repeats. Deletions are indicated by crossed lines, insertions are shown in yellow, and BRC repeats are shown in red. Deletion and insertion sizes and genomic coordinates are indicated for each clone. Clones shown in red contain genomic events that restore the BRCA2 reading frame at the DNA level. Clones shown in black either disrupt the canonical exon 11 splice donor or retain an out-of-frame genomic sequence and are therefore predicted to restore BRCA2 expression through splicing-mediated restoration. (B) Functional re-targeting of representative p.Y1569Rfs*2 reversion clone 1 and p.S1982Rfs*22 reversion clone 6. Each clone was edited with an sgRNA targeting *BRCA2* exon 3 upstream of the exon 11 restoration event, generating unedited, secondary frameshift, and in-frame alleles. Bars show the mean log2 fold change in allele frequency following mock or 1 μM niraparib treatment relative to day 7. Individual biological replicates are shown as dots. Statistical significance is indicated as follows: NS, not significant; *$P < 0.05$, **$P < 0.01$; ***$P < 0.001$.

#### Horacek et al. Supplemental Figure 8

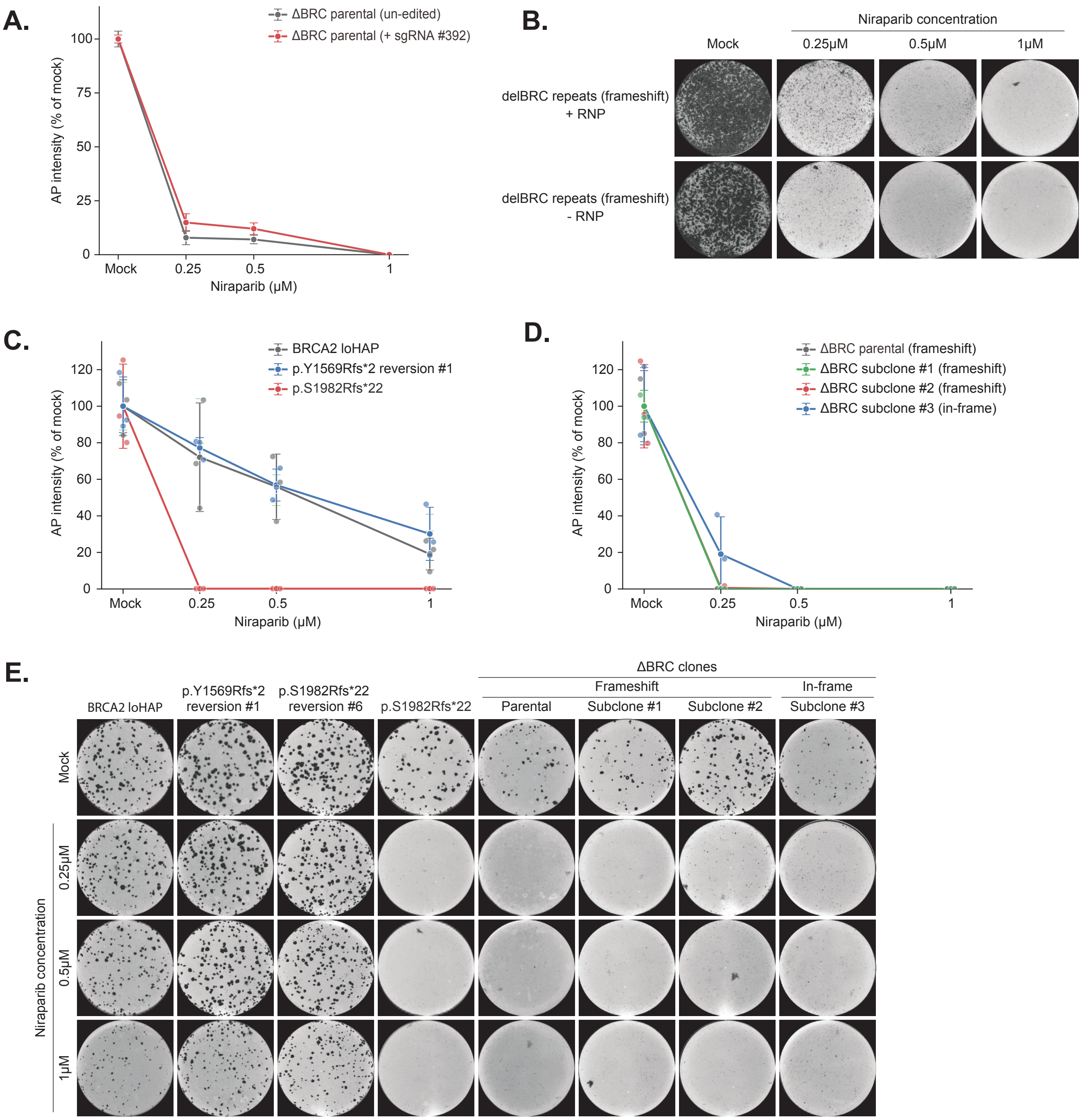

**Supplemental Figure 8: *BRCA2* restorations lacking BRC repeats fail to confer PARPi resistance.** (A) Quantification of alkaline phosphatase (AP) staining in the ΔBRC parental frameshift line with or without re-editing by sgRNA #392 following treatment with mock media or the indicated concentrations of niraparib. Three biological replicates per condition. AP intensity was normalized to the corresponding mock-treated condition. Dots represent mean normalized AP intensity, while lines represent the standard deviation across replicates. (B) Representative AP-stained wells corresponding to the experiment in (A). (C) Quantification of AP staining in unedited *BRCA2* loHAP cells, p.Y1569Rfs*2 reversion clone 1, and the parental p.S1982Rfs*22 frameshift line following niraparib treatment. (D) Quantification of AP staining in the ΔBRC parental frameshift line, two compound frameshift ΔBRC subclones, and the in-frame ΔBRC subclone #3 following niraparib treatment. (E) Representative AP-stained wells for the indicated parental, reversion, and ΔBRC genotypes following treatment with mock medium or the indicated concentrations of niraparib. Three replicates were stained per condition. Dots represent mean normalized AP intensity, while lines represent the standard deviation across replicates.
